# ProcessNets: Towards an efficient approach for ensemble analysis of biological networks

**DOI:** 10.64898/2026.09.22.753520

**Authors:** Sugyani Mahapatra, Manikandan Narayanan

## Abstract

Integrative analysis of the properties of multiple networks pertaining to a biological system is a key problem in systems biology. But computing the properties of thousands of large networks poses a challenge. Most techniques that partially address this challenge, such as parallelism and optimized algorithms, treat the analysis of each network as an independent task. However, the networks arising from the same biological system are often similar to each other, and can we harness this similarity to achieve speedup? Towards this end, we propose a phylogeny-like data structure, `nc`-tree, to compactly represent an ensemble of related biological networks; and a versatile frame-work, ProcessNets, that runs incremental algorithms over the `nc`-tree to speedup property computations on the input networks. Our asymptotic analysis of space and time complexity, coupled with empirical evaluations of diverse ensembles of simulated and real-world GTEx gene coexpression networks, demonstrate that ProcessNets leads to significant space and time advantages when computing different network science measures on ensembles of similar networks. For instance, we achieve a compression factor of 2.1*×* and 8.1*×* in storing two representative ensembles of GTEx Muscle Skeletal networks with 1000 nodes/genes, and a time speedup of 2.86*×* to 3.97*×* over the fastest baseline for computing degree centrality in ensembles of networks with at least 14, 500 nodes derived from subsampled GTEx datasets of ten tissues. These results are promising and encourage application of ProcessNets to analyze biological network ensembles derived from rapidly accumulating consortium/biobank-based datasets.

## I. Introduction

Omics datasets have been represented as networks, wherein biological entities such as genes, proteins, metabolites, drugs, and diseases are represented as nodes, and the underlying functional, physical, or chemical interactions among these entities are represented as edges [1], [2]. A modern omics study rarely involves a single network though; instead it often has to consider multiple networks derived from different omics layers, biological conditions, computational analyses, statistical resamplings, or patient groups. Integrative analysis of an ensemble of such networks provides a more comprehensive view of the underlying system. Among various integration strategies, the “late integration” strategy analyzes each network independently, and combines the resulting outputs such as modules, scorings, rankings or other relevant network statistics at a later stage [3]. Late integration thus preserves network-specific information and enables analysis of how these outputs vary across the networks [4].

The authors are with Department of Computer Science & Engineering, Centre for Integrative Biology and Systems medicinE, and Wadhwani School of Data Science & AI, Indian Institute of Technology (IIT) Madras, Chennai, India. Second author is also affiliated to Sudha Gopalakrishnan Brain Centre, IIT Madras, Chennai, India.

To realize late integration, we need efficient computational methods that can integrate say hundreds to thousands of large biological networks derived from a consortium-based or biobank-scale omics dataset. Consortium-based efforts such as The Genotype-Tissue Expression (GTEx) [5] and The Cancer Genome Atlas [6] have routinely used high-throughput technologies like RNA sequencing to profile all human genes, including ≈19, 000 protein-coding genes, across several hundred donors; and the UK Biobank study [7] has quanti-fied ≈ 3, 000 proteins across ≈ 54, 000 participants. Many biological networks derived from these datasets, particularly those constructed using correlation or other similarity-based metrics, can be highly dense. High edge density renders most graph-theoretic algorithms pertaining to centrality computation, spectral clustering, and community detection computationally expensive [8], [9]. Although parallel and high-performance computing platforms can partially alleviate this computational bottleneck, the scalability of this approach to handle massive network ensembles is still constrained by available computational resources.

Existing studies primarily address the computational challenges through feature selection and dimensionality reduction [3], which can potentially discard relevant topological information. Alternatively, studies have focused on improving integration methodologies [1], or accelerating graph-theoretic algorithms for network analysis [9], [10]. But these approaches continue to treat the analysis of each network as an independent computational task, despite the networks in an ensemble being derived from the same dataset/source and hence similar to each other. This leaves open the question of whether the similarity of networks in an ensemble can be harnessed to speedup the computational tasks on all input networks. In other words, despite the widespread adoption of late integration, the problem of efficiently computing a network science measure across an ensemble of multiple similar biological networks remains largely unexplored.

To address the above problem, this study proposes a versatile framework, ProcessNets, that introduces a compact representation of the networks in an input ensemble and uses it to repeatedly invoke an incremental dynamic graph algorithm. More specifically, our proposed framework exploits the structural similarity among the networks to hierarchically organize them into a reusable tree data structure, hereafter referred to as the Networks’ Cascade Tree (`nc`-tree); this tree resembles a phylogeny-like evolutionary structure, wherein the root graph progressively evolves into the network at each tip via a sequence of edge increments. Under this paradigm, every input network in the ensemble is interpreted as a dynamic graph [11] that originates from the root and systematically evolves through a sequence of incremental updates (batch edge insertions) along the branches of the `nc`-tree.

This `nc`-tree paradigm leads to immediate space and time advantages for an ensemble of similar networks. Instead of storing each network separately, we can store all networks compactly using only the `nc`-tree, including the root network and incremental edges along the branches. Similarly, instead of computing the measures of each network separately using traditional (static) algorithms, we can update the measures efficiently along each branch of the `nc`-tree using incremental dynamic graph algorithms. Incremental dynamic algorithms often use intermediate data structures (DS) to update measures efficiently upon edge insertions, which are required between the successive graph states along a branch [11]. In our framework, we prefer incremental over fully dynamic algorithms, as the latter incurs more overhead in handling both edge insertions and deletions.

The key contributions of our study are three-fold:

- *Methodological:* Given an ensemble of similar yet distinct networks, defined over the same set of nodes, we present the ProcessNets framework to efficiently compute different global/local properties of each network. As part of this framework, we design `nc`-tree, the phylogeny-like data structure that compactly stores the networks in the ensemble.
- *Algorithmic:* We compare the asymptotic time complexity of ProcessNets to the trivial/baseline approach and discuss different factors that influence our framework’s speedup. Additionally, we adapt an existing Steiner tree algorithm to develop a scalable heuristic for constructing the `nc`-tree.
- *Empirical:* We measured the running times of different configurations of our framework vs. the baseline for computing three network science measures in simulated or real-world (GTEx) network ensembles. We carefully interpret these results and provide recommendations on which approach to use based on the characteristics of the input ensemble to achieve the best speedup.

The promising results of our versatile framework on various simulated and real-world ensembles should encourage applica-tion of ProcessNets to analyze biological network ensembles derived from rapidly accumulating consortium/biobank-based datasets.

## II Methods

### A. Problem Statement and Framework Outline

Given an ensemble of *B* graphs (referred to also as networks), each defined on the same set of nodes *V*, but with different edge sets (*E*_i_ for the *i*^th^ graph *G*_i_) and represented as ℊ = {*G*_1_ = (*V, E*_1_), *G*_2_ = (*V, E*_2_), …, *G*_B_ = (*V, E*_B_)}, we address the problem of computing a network science measure/property *P*, defined either for every node or at the global scale, in each of these networks (denoted as *P* (*G*_1_), *P* (*G*_2_), …, *P* (*G*_B_)).

A trivial/baseline approach to solve this problem is to iteratively compute the measure *P* in each of the networks independently using any appropriate static algorithm. However, this approach can be computationally expensive in large-scale ensembles and for complex measures with high asymptotic complexities. Our proposed framework, “PROperty Computation in EnSembleS of NETworkS (ProcessNets)”, exploits the similarity among the edge sets of the networks to organize them into a “rooted tree structure”, namely, the `nc`-tree, and hierarchically computes the measure *P* for all networks in the ensemble through repeated invocations of an incremental dynamic graph algorithm (Fig. 1). Though our framework works correctly for any collection of input networks, it is designed to exploit similarities among the networks.

**Fig. 1.**
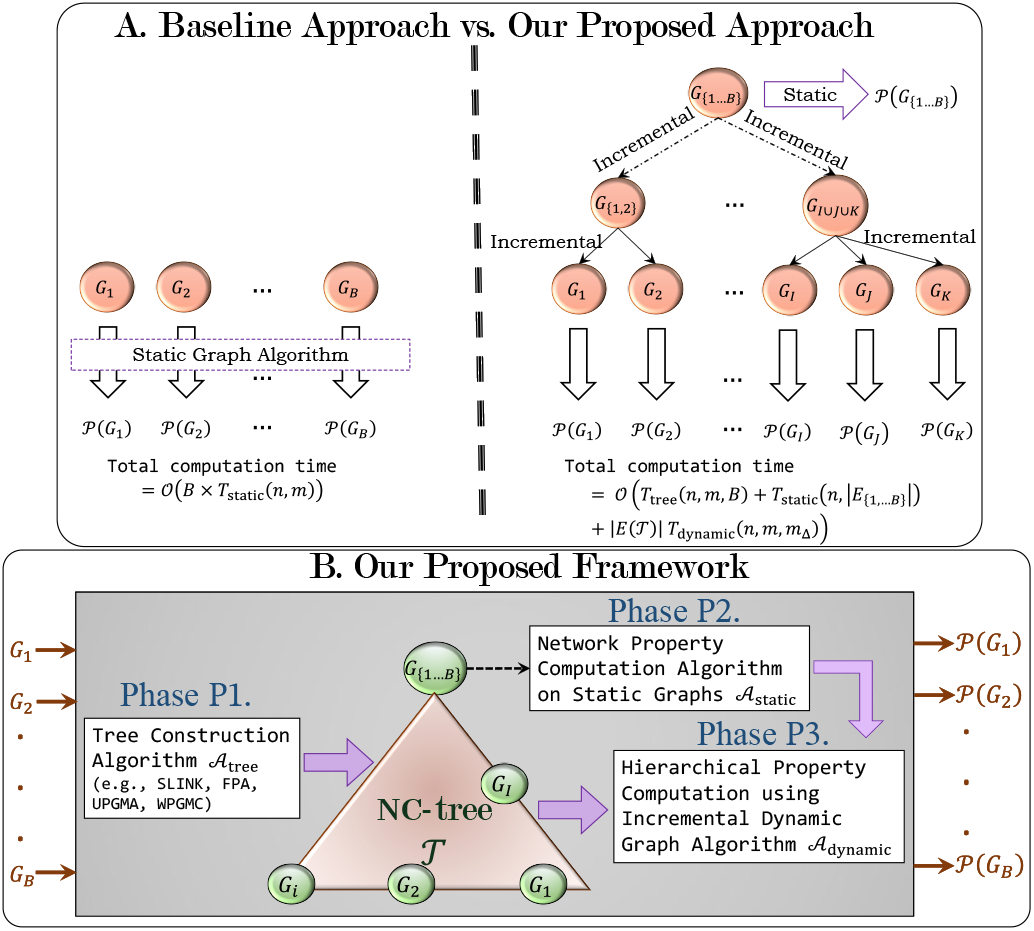
Overview of the ProcessNets Framework. A. Comparative overview between the baseline approach and our framework, along with their asymptotic time complexities. Here, *T*_tree_, *T*_static_ and *T*_dynamic_ represent the time complexities of *A*_tree_, *A*_static_ and *A*_dynamic_ respectively. *G*_*{*1,2*}*_ (parent of *G*_1_ and *G*_2_) and *G*_*I∪J*_ (parent of any networks *G*_*I*_ and *G*_*J*_ in *T*) represent examples of a few possible intermediate branching points in T. B. The outline of the framework depicting three algorithms for its three phases.

To quantify this similarity, we use two characteristic feature of the ensemble, namely, “`MPJS`” and “average edge count (EC)”. The structural similarity between two graphs *G*_i_ and *G*_j_ can be quantified using the Jaccard Similarity Index `JSI` 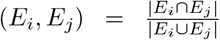. The overall similarity of networks in the ensemble *ℊ* can then be quantified using the Mean Pairwise Jaccard Similarity (`MPJS`), estimated from a randomly chosen subset of 10 networks as 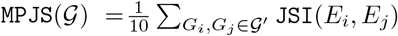, where ℊ^′^ ⊆ *ℊ* such that | *ℊ*^′^ | = | *ℊ* | ≥ 10 (assuming 10). Average edge count (or average EC) denotes the average number of edges in the networks of the ensemble, 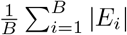.

### B. Description and Analysis of the Framework

We now describe our ProcessNets framework in detail, including description of `nc`-tree, different phases of the framework, and its correctness and running time analysis.

1. <u>*The Networks’ Cascade Tree*:</u> The `nc`-tree is defined as an out-directed rooted tree *T* = (*V* (*T*), *E*(*T*)) such that there is a one-to-one correspondence between the tips (or leaves) of the tree and the networks in the input ensemble *ℊ*. Additionally, each branching point (internal node) of *T* maps to a graph *G*_I_ whose edge set is the edges shared among every network in the tips of the clade (subtree) rooted at this branching point. Formally, let *J* ⊆ {1, 2, …, *B*} and *E*_J_ =⋂_j∈J_ *E*_j_. Then, we can define *G*_I_ = (*V, E*_I_), where *I* = {*i* ∈ {1, 2, …, *B*} | *G*_i_ *is in a tip of the clade rooted at the branching point corresponding to G*_I_} (see Fig. 1A for examples). This ensures that new edges can only be inserted but not deleted between nodes while traversing from a *parent network* to a *child network* along a branch of the nc-tree. It may be noted here that if a branching point and its child are the same, then there may be no edge additions. Moreover, a branching point with exactly one child may also be compressed with its parent, ensuring that there are no single-child branching points. Graph edges that are added to a parent network to construct its child network along a branch of `nc`-tree are hereafter referred to as “incremental edges”, and we define the cost of the `nc`-tree, *T* as the total number of incremental edges across all its branches. Given an ensemble of networks, *G*, the optimization problem of constructing *T* with minimum cost is NP-hard, because it is equivalent to the Camin-Sokal phylogeny construction problem [12], [13] (using a row-major, bit-vector representation of the adjacency matrices of the networks in). Despite the existence of several parameterized and approximation algorithms [14] to address this problem, their running times make them impractical for ensembles on large-scale biomolecular networks. Thus, we focus on constructing valid `nc`-trees (even though they may not be optimal) using scalable heuristics.
2. <u>*The Three Phases*:</u> The ProcessNets framework runs in three phases (Fig. 1B) as described below.

### P1. nc-tree Construction

Given the input ensemble of networks, the framework uses heuristic algorithms such as the `rstar` or agglom-erative clustering algorithms to construct the `nc`-tree *T*. Detailed descriptions of these algorithms are provided in Section II-C. We use *A*_tree_ to denote this tree construction algorithm. It may be noted here that the constructed `nc`-tree is reusable in computation of multiple network science measures; consequently, this phase of the framework need not be repeated for a fixed ensemble.

### P2. Measurements of the Root Graph P (G_{1,…,B}_)

Similar to that in the baseline approach, an appropriate algorithm (*A*_static_) to compute *P* in (static) graphs is used to compute the measure in the root *G*_{1,…,B}_ = (*V, E*_{1,…,B}_).

### P3. Hierarchical Computation at Branching Points

Finally, the framework uses an (edge-addition only) incremental algorithm for dynamic graphs (A_dynamic_) to compute the measure at every branching point (and tip) in the `nc`-tree hierarchically in a top-down approach. The algorithm is applied once for every branch of the `nc`-tree, using a level-order traversal of the tree starting from the root, and uses the measurements of the parent network to compute the same in the child.

We redirect the readers to Appendix A.A and Algorithm S1 for a summary of the framework and its pseudocode respectively.

### 3. Correctness

Given that the `nc`-tree is valid, i.e., every network in the ensemble is in *T* and every child network is formed by inserting edges to its parent network, for each network *G* ∈ ℊ, P (*G*) can be correctly computed by:(i) computing *P* of the root network *G*_{1,…,B}_ using *A*_static_, and (ii) by applying *A*_dynamic_ repeatedly along the edges of the path from the root to the tip in corresponding to *G*. This intuitive proof can be formalized using mathematical induction. Thus, the measurement for every network in the ensemble can be correctly computed with a tree traversal operation.

### 4. Asymptotic Efficiency

We use the notations *n* and *m* to represent the number of nodes in any network in the ensemble and the maximum number of edges in a network of the ensemble respectively, i.e., *n* = |*V* | and m = max_*i∈*{1,…,*B*}_ |*E*_*i*_|.

#### Time Complexity

Three algorithms used in the three phases of the framework constitute three components that directly affect the asymptotic time complexity of our framework. Clearly, running time of the `nc`-tree construction *algorithm A*_*tree*_ depends on the size and number of input graphs, and we use *T*_tree_(*n, m, B*) to represent its time complexity. In the second phase, the static algorithm *A*_static_ is applied only to the root graph, *G*_{1,…,B}_. Asymptotic time complexity of static algorithms usually depend on size of the input network, and hence we represent the time complexity of this algorithm as *T*_static_(*n*, |*E*_{1,…,B}_|).

Apart from the new set of incremental edges that are added to a parent network, dynamic incremental algorithms *A*_dynamic_ generally require intermediate data structures (DS) that compactly represent structural information (about the parent network) to compute the measure. Sizes of these intermediate DS are bounded by the size of the graph and thus, we use *T*_dynamic_(*n, m, m*_Δ_) to represent the worst-case time complexity of *A*_dynamic_; here *m*_Δ_ represents the maximum number of edges added to a network along a branch of the `nc`-tree, i.e., *m*_Δ_ = max_e=(*G*_parent_,*G*_child_)∈E(T)_ |*E*_child_ \*E*_parent_|. Now, this algorithm is applied once for every branch of the `nc`-tree, and hence, contributes a factor of *O* (|*E*(*T*)| *T*_dynamic_(*n, m, m*_Δ_)) to the total time complexity.

Thus, the worst-case asymptotic time complexity of the framework is (*T*_tree_(*n, m, B*) + *T*_static_(*n*, | *E*_{1,…,B}_ |) + *E*(*T*) *T*_dynamic_(*n, m, m*_Δ_)) (Fig. 1A).

It may be noted here that in a *k*-ary tree, i.e., the maximum number of children at any branching point of the tree is at most *k*, |*E*(*T*) |= *O*(*kB*). Usually, since *k* is fixed and small, the number of branches in the tree can be bounded as |*E*(*T*) | = *O*(*B*), and thus, the time complexity of the framework can be represented as *O*(*T*_tree_(*n, m, B*) + *T*_static_(*n*, | *E*_{1,…,B}_ |) + *B T*_dynamic_(*n, m, m*_Δ_)).

Comparing this asymptotic time complexity with the baseline approach’s *O*(*BT*_static_(*n, m*)) time (see Fig. 1A), the ProcessNets framework gains an advantage under the following scenarios:

- Either a time-efficient algorithm is used for constructing the `nc`-tree or an existing `nc`-tree is being reused.
- The networks in the ensemble are very similar to each other (i.e., the ensemble has high `MPJS`) enabling many edges in the root graph and large number of incremental edges in branches closer to the root.
- The incremental dynamic graph algorithm used is highly efficient and runs in time faster than the static graph algorithm.

#### Space Complexity

To represent the input ensemble using an `nc`-tree, we need to store only the edge set of the root and information pertaining to each branch (i.e., IDs of the parent and child network and the set of incremental edges that are added to the parent in order to obtain the child). Thus, the space complexity of such an `nc`-tree is bounded by *O* (| *E*_{1,…,B}_ | + *E*(*T*) *m*_Δ_). Similar to that for time complexity, in a *k*-ary `nc`-tree with fixed and small values of *k*, |*E*(*T*) | = *O* (*B*), and hence, the space required is (*E*_{1,…,B}_ + *Bm*_Δ_). Conversely, the baseline representation of the networks as adjacency lists require *O* (Σ_i∈{1,…,B}_ |*E*_i_|) = (*Bm*). It may be noted here that if the networks are similar, i.e., the ensemble has a high `MPJS`, then *m*_Δ_ *<<* min_i∈{1,…,B}_ | *E*_i_|.

Thus, the `nc`-tree provides a compact and space-efficient representation of the input ensemble. Additionally, the internal cascading structure enables full reconstruction of the tree (along with the input ensemble) with a simple tree traversal algorithm.

### C. NC*-tree Construction Algorithms/Heuristics*

We propose a heuristic algorithm to construct the `nc`-tree and discuss other algorithms that were used to construct different variants of the `nc`-tree for empirical analyses.

#### 1) The rstar Algorithm

As mentioned above, the optimal `nc`-tree construction problem is NP-complete. We, therefore, adapt an existing parameterized-approximation algorithm using the parameter *p* (i.e., the number of Steiner nodes) in [14] into a heuristic, hereafter named as “*Recurrent Star* (`rstar`)” algorithm, to construct a variant of the `nc`-tree*T*_rstar_. The underlying idea of our proposed algorithm is to iteratively group networks with the highest number of edges and replace them with a representative network, constructed by intersecting the edge sets of all networks in this group; branches are then added from this representative network to every network in the group. However, if there is only one network in the group (during any iteration), then this network is grouped along with other networks consisting of the second-highest number of edges, and this newly-constructed group is considered for replacement.

This heuristic has been summarized in Algorithm 1. Clearly, for every network *G* ∈ ℊ, *level*[*G*] ≠ 0, and hence *G* is grouped and forms exactly one branching point/tip of *T*_rstar_. Also, since the parent node is constructed by intersecting the edge sets in the corresponding group, every child network can be formed by only inserting edges to its parents. Thus, the resulting tree *T*_rstar_ is a valid `nc`-tree.

##### Algorithm 1

Recurrent Star (`rstar`) algorithm.

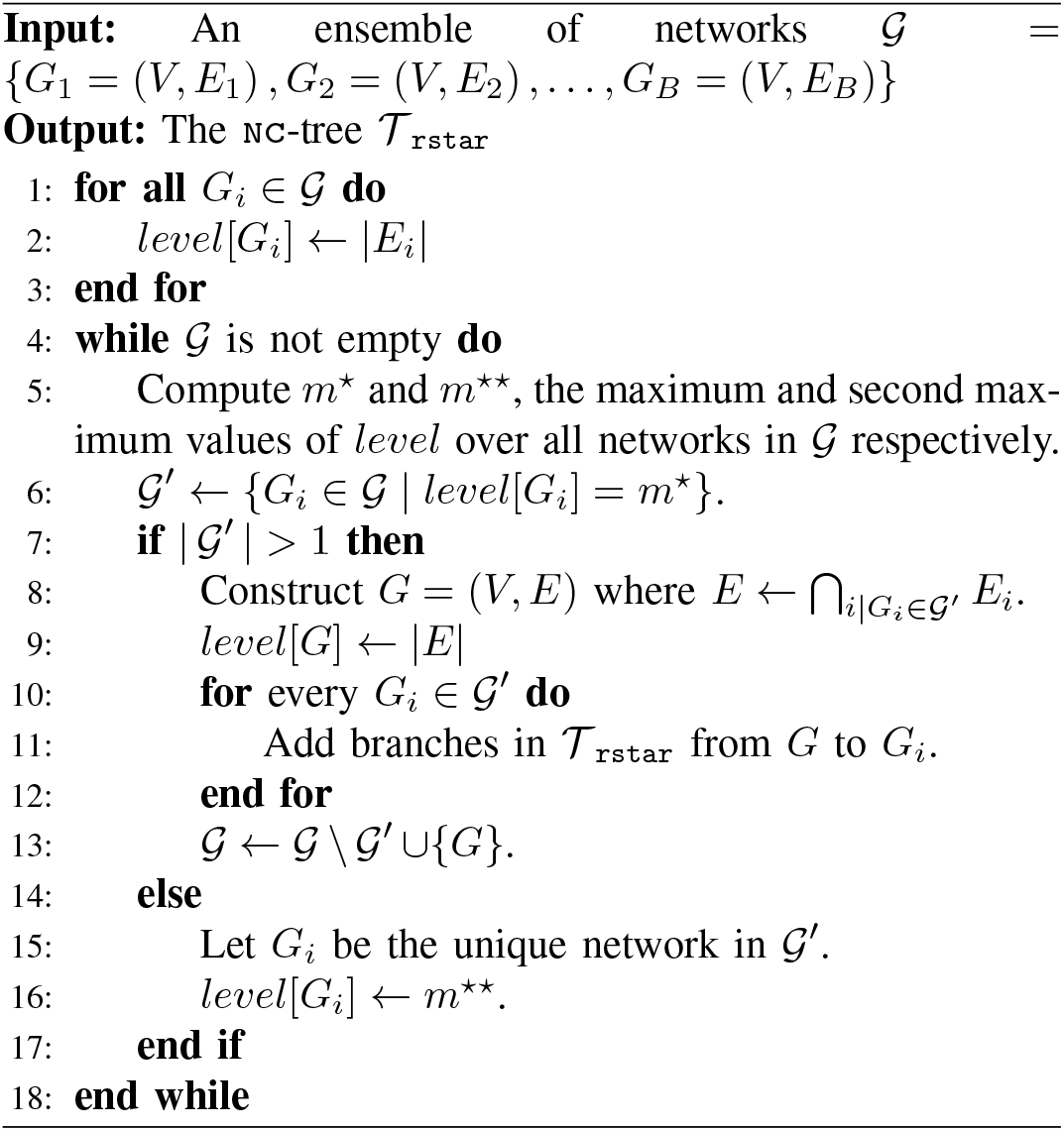

An implementation of the *level* list using max-heaps yields an asymptotic running time of *O*(*Bm* log *B*) for the proposed `rstar` algorithm. However, for our empirical analysis, we use a simpler array-based implementation; although this variant exhibits *O*(*B*^2^*m*) time complexity, it is still faster than other `nc`-tree construction algorithms (cf. Section IV-A).

#### 2) Clustering-Based Algorithms

Apart from this, we used four different hierarchical (agglomerative) clustering algorithms, namely, single-linkage (SLINK), complete-linkage or farthest point algorithm (FPA), group average-linkage (UPGMA), and median-linkage (WPGMC) [15], to generate *T*_slink_, *T*_fpa_, *T*_upgma_ and *T*_wpgmc_ respectively. These hierarchi-cal clustering algorithms start with each graph in the ensemble as an individual cluster and iteratively merge a pair of similar clusters into a cluster at the next level of hierarchy. This process results in a hierarchical organization of the networks in the ensemble that brings together graphs with similar edge sets into clusters at different levels.

Pair of clusters with the minimum inter-cluster distance is merged at any iteration. We generated minhash signatures [16] of length 10 for each network and used the Hamming distance between the signatures of every pair of network as the initial distance metric for these algorithms (see Appendix A.B for details). Following the formation of each cluster, the distance of the cluster to other clusters is determined based on the algorithm – for instance, in the SLINK algorithm, distance of the newly-formed cluster to another existing cluster is determined as the minimum distance between any pair of their networks (see [15] for similar details on other algorithms).

### D. Configurations of ProcessNets and Baseline Approach

The ProcessNets framework was implemented us-ing the five different algorithms to construct the `nc`-tree as described above, resulting in five configurations of the framework implementation, namely, ProcessNets-rstar, ProcessNets-slink, ProcessNets-fpa, ProcessNets-upgma and ProcessNets-wpgmc. The baseline approach mentioned above (see Section II-A) was implemented using two static algorithm variants— (i) self implementation (de-noted `baseline`-self) and (ii) using library functions from NetworkX package (denoted `baseline`-networkx). We collectively refer `baseline`-self and `baseline`-networkx as the “`baseline`-* configurations” or simply “`baseline`-*”; similarly we collectively refer the five configurations of ProcessNets as the “ProcessNets-* configurations” or sim-ply “ProcessNets-*”. We measured the running time taken by these ProcessNets-* and `baseline`-* configurations to compute four network science measures on different ensembles of networks, and report the performance trends in Results section.

### E. Algorithms for Network Science Measures

The efficiency of ProcessNets framework is influenced by the efficiency of the incremental algorithms (as discussed in Section II-B4). Recall that the incremental graph algorithm is run once along each branch of the `nc`-tree, in order to process the batch of edges added when traversing from the parent to the child network of the branch. We prefer incremental (dynamic) graph algorithms with the following two characteristics to improve the running time of our framework.

– *Use Intermediate DS only?* We prefer incremental algorithms that compute the updated measurements of the child network using only an intermediate DS with limited size, without needing access to the larger parent network DS.]
– *Process Edge Insertions as a Batch?* We prefer algorithms that process all incremental edges from the parent to the child network as a single batch, rather than adding them sequentially one edge at a time.

Based on whether an incremental algorithm satisfies the above two preferences, we’ve four combinations and chose to analyze four corresponding network science measures (Table I): degree centrality, component size (defined as size of the connected component containing a particular node [1]), PageRank (PR) centrality, and closeness centrality. A brief summary of these measures along with the algorithms used to compute them are given in Table I and Appendix A.C. Since the incremental algorithm for closeness centrality doesn’t satisfy the above two characteristics, we only perform certain preliminary analyses to verify the subpar performance of the framework for closeness centrality (see Appendix B.A and Fig. S1), and focus the rest of the manuscript on the other three measures.

**TABLE I.**
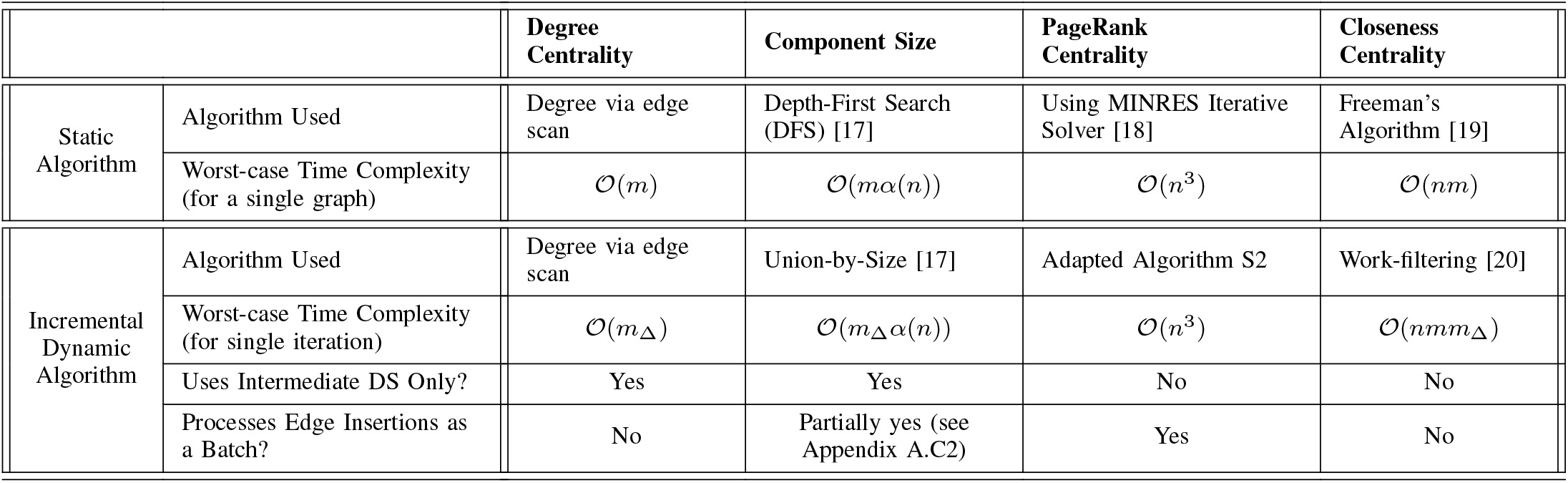
Summary of algorithms focused in this work for computing network science measures on an undirected, unweighted graph with *n* nodes and *m* edges. Also, *m*_Δ_ denotes the number of edges inserted in one batch update of an incremental algorithm into the original graph to obtain the final graph with *m* edges, and *α*(*n*) denotes the inverse Ackermann function.

|  |  | Degree Centrality | Component Size | PageRank Centrality | Closeness Centrality |
| --- | --- | --- | --- | --- | --- |
| Static Algorithm | Algorithm Used | Degree via edge scan | Depth-First Search (DFS) [17] | Using MINRES Iterative Solver [18] | Freeman's Algorithm [19] |
| | Worst-case Time Complexity (for a single graph) | $\mathcal{O}(m)$ | $\mathcal{O}(m\alpha(n))$ | $\mathcal{O}(n^3)$ | $\mathcal{O}(nm)$ |
| Incremental Dynamic Algorithm | Algorithm Used | Degree via edge scan | Union-by-Size [17] | Adapted Algorithm S2 | Work-filtering [20] |
| | Worst-case Time Complexity (for single iteration) | $\mathcal{O}(m_\Delta)$ | $\mathcal{O}(m_\Delta\alpha(n))$ | $\mathcal{O}(n^3)$ | $\mathcal{O}(nmm_\Delta)$ |
|  | Uses Intermediate DS Only? | Yes | Yes | No | No |
|  | Processes Edge Insertions as a Batch? | No | Partially yes (see Appendix A.C2) | Yes | No |

### F. Empirical Analysis: Setup, Datasets, and Ensembles

#### 1) Setup

We constructed several ensembles of simulated and real-world networks with varied `MPJS` to empirically analyze the performance of different configurations of our proposed framework with that of the baseline approach. We performed multi-factor scalability analysis to evaluate the speedup achieved by our framework over the baseline approach. We primarily focus on (undirected and unweighted) gene-gene co-expression networks constructed on bootstrapped/subsampled gene expression datasets (generated by sampling from a reference transcriptomics dataset using bootstrapping/subsampling respectively). Details on the construction of a coexpression network from a single bootstrapped/subsampled dataset is in Appendix A.D. Details on the implementation of configurations for computing network science measures in all constructed ensembles is in Appendix A.E (see also Table S1 for how dedicated cores were used to benchmark each configuration).

In this study, our primary goal is to compare the total running time of the ProcessNets-* and `baseline`-* in computing the measures. This total time includes the time taken to read the input ensemble from files, the processing time taken to compute the measure, and the time taken to write the output. Note that the input for the `baseline`-* is the actual networks in the ensemble, whereas the input for the ProcessNets-* is the corresponding `nc`-tree. Since `nc`-trees are reusable, we treat `nc`-tree construction as a preprocessing step and consider the precomputed `nc`-tree as input for the ProcessNets-*.

#### 2) Simulated Network Ensembles

##### a) Simulated Ensemble I (Comprising Similar Networks)

We used a similar formulation as in [21] to simulate a population gene expression dataset with *n* genes (categorized into two sets) and one million samples. Gene expression values for the first set of *n/*2 genes were generated by sampling from a standard normal distribution, whereas the gene expression values for the second set of *n/*2 genes were generated as a linear combination of the expression values of a randomly chosen set of neighbors from the former set. From this population, an observed dataset with *s* samples was drawn by sampling uniformly at random without replacement. Bootstrapping was performed on this observed dataset by drawing *s* samples uniformly at random with replacement, and a coexpression network was constructed on each bootstrapped dataset; this (bootstrapping) process was repeated *B* times to generate an ensemble of *B* bootstrapped coexpression networks.

We varied the parameters *n, B* and *s* to generate multiple ensembles and evaluated the performance of both the framework and the baseline approach across these settings. More specifically, we performed two distinct analyses:

- (*n, B*)*-effect with fixed s*: We varied *n* and *B* (both from 100 to 1000) with 500 samples in the observed dataset to assess the effect of size of the ensemble on the running time. For each ensemble with a specific value for both *n* and *B*, we computed the three considered network science measures, namely, degree centrality, component size, and PR centrality, using all seven configurations and compared their run times with scatter plots. `MPJS` of these ensembles range between 0.72 and 0.81.
- (*n, s*)*-effect with fixed B*: Here, we fixed *B* at 1000 and varied *n* (from 100 to 1000) and *s* (over selected values in the range [75, 1500]). We performed similar analyses as above for all the three measures using the seven configurations. `MPJS` of these ensembles range in [0.63, 0.92], and although `MPJS` increases with *s*, the rate of increase slows down for large *n*.

##### B) Simulated Ensemble II (Comprising Dissimilar Networks)

As discussed above, similarity among the networks in the input ensemble is a key factor influencing the performance of our proposed framework (cf. Section II-B4). To empirically demonstrate its effect on the running time efficiency of the framework relative to the baseline approach, we simulated an ensemble of dissimilar networks and showed that the ProcessNets-* take longer time to compute the same network science measures than the `baseline`-* under similar conditions as in Simulated Ensemble I.

We sampled a degree sequence from a power-law distribution with the exponent parameter *γ* = 2.5 and generated a network consistent with this degree sequence; this process was repeated independently 1000 times to construct an ensemble of *B* = 1000 networks and `MPJS` ≈ 0.1. We varied *n* from 100 to 1000 to generate multiple such ensembles and performed similar analysis on the running times to compute the three network science measures by all configurations as in Simulated Ensemble I.

#### 3) Real-World Network Ensembles

We used gene expression values of protein-coding genes for 10 tissues, profiled across varied samples, from their single-tissue cis-expression Quantitative Trait Locus (eQTL) dataset provided by the GTEx consortium [5] Release V8 (downloaded on February 2025 from the GTEx portal) to construct several ensembles. Filtering of samples and genes, and preprocessing (including covariates’ adjustment) of these datasets were done as per GTEx recommended practices following our earlier study [21]. These filtered and preprocessed datasets are used for inferring the gene-gene coexpression networks.

For any given GTEx tissue, we can construct three categories of ensembles based on the resampling method applied, and the fraction of all *s* samples in the gene expression dataset that are resampled.

*ℬs. By bootstrapping from all the samples*: Generate a bootstrap resampled dataset by sampling with replacement *s* samples from the original dataset, and use it to construct a gene-gene coexpression network. Repeating this process 1000 times generates an ensemble with *B* = 1000 networks. Now, depending on which genes are used to construct the ensemble, this analysis can be performed in two variants:

i. using all genes filtered for the tissue (i.e., *n* ≈ 15000 genes; these ensembles have `MPJS` ∈ [0.2, 0.24] across the tissues considered), or
ii. using only *n* top-varying genes (i.e., for each of the ten values of *n* considered between 100 and 1000, an ensemble is constructed on a smaller dataset containing only the top-*n* genes with the highest variance across all samples).

*Ss. By subsampling* 90% *of the samples:* Construction of ensembles in this category also involves a similar process as in the previous category (i.e., category *ℬs*); however, instead of bootstrapping all samples, only *f* = 90% of the actual samples is sampled without replacement to obtain a subsampled dataset. Again similar to the previous category, we can generate two variants of ensembles, depending on the genes chosen to construct the ensemble:

i. using all genes filtered for the tissue (`MPJS` ∈ [0.55, 0.61]), or
ii. using only *n* top-varying genes.

*Ssf*. *By subsampling f* % *of the samples*: Finally, this category is similar to category *Ss*.(i), except that, instead of fixing the subsampling fraction (*f*) at 90%, we construct an ensemble for each of the six values of *f* ranging from 70% to 95%.

We generated ensembles in each of the above categories for the GTEx tissue with the largest sample size, viz., Muscle Skeletal tissue. These ensembles exhibited varying degrees of network similarity (e.g., range of MPJS being [0.44, 0.58] for category *ℬs*.(ii) ensembles, [0.86, 0.92] for *S s*.(ii) ensembles, and [0.44, 0.69] for *sf* ensembles). For every other tissue considered, we generated ensembles only from categories *ℬs* and *Ss*, and further focused only on variant (i), i.e., using all genes filtered for the tissue (see Table II). An example of outputs generated on the ensembles constructed using the Whole Blood tissue is given in Supplementary Data S1.

**TABLE II.** Tissues whose gene expression datasets (from gtex [5] v8) were used to construct ensembles of networks in this study. Here, *n* refers to the number of filtered protein-coding genes and *s* is the sample size of the dataset. Average edge count is rounded to the nearest integer.

| Tissue | $n$ | $s$ | Average edge count of an ensemble in | |
| --- | --- | --- | --- | --- |
| | | | Category $\mathcal{B}s$<br>MPJS $\in [0.22, 0.24]$ | Category $\mathcal{S}s$<br>MPJS $\in [0.59, 0.61]$ |
| Muscle Skeletal | 15163 | 706 | 3,133,882 | 219,307 |
| Whole Blood | 14582 | 670 | 2,360,759 | 131,921 |
| Skin Sun Exposed (Lower Leg) | 16646 | 605 | 3,489,116 | 204,052 |
| Thyroid | 16607 | 574 | 3,307,366 | 197,632 |
| Lung | 16765 | 515 | 3,326,089 | 162,468 |

(b) Tissues with moderate sample size
| Tissue | $n$ | $s$ | Average edge count of an ensemble in | |
| --- | --- | --- | --- | --- |
| | | | Category $\mathcal{B}s$<br>MPJS $\in [0.2, 0.22]$ | Category $\mathcal{S}s$<br>MPJS $\in [0.55, 0.59]$ |
| Stomach | 16485 | 324 | 2,721,483 | 76,787 |
| Pancreas | 15848 | 305 | 2,172,754 | 41,812 |
| Pituitary | 17036 | 237 | 3,553,676 | 129,170 |
| Brain Cerebellum | 16463 | 209 | 2,669,982 | 66,477 |
| Brain Cortex | 16536 | 205 | 2,833,775 | 72,751 |

## III. Results

We set out to examine if the asymptotic efficiency of our ProcessNets framework, relative to the baseline approach for computing a property of each network in an ensemble, actually translates into practice. Towards this end, we compared the running times of the ProcessNets-^*^ vs. baseline-^*^ on sev-eral simulated and real-world ensembles with varying degrees of network similarity and average edge count.

### A. Results on Simulated Ensemble I (Comprising Similar Networks)

In an ensemble of similar networks (Simulated Ensemble I), ProcessNets-^*^ consistently show significant improvements in running times than the baseline-^*^ (see Fig. 2A-B and “*S2*.*1 Simulated Ensemble I*” in Supplementary Data S2). The magnitude of this improvement, i.e., the speedup, increases with increase in values of the three factors, i.e., *n, B* and *s* (Fig. 2A-B). For instance, in degree centrality computation, the speedup (on the average running time across the three trials) of ProcessNets-rstar over the baseline-networkx configuration ranges between 1.22× and 2.84× when *n* = 100 and between 3.88 × and 6.46 × when *n* = 1000 for varied *s*. Similarly, for PR centrality, the speedup of ProcessNets-slink over the baseline-self configuration increases from 0.72 × to 1.27 × and from 7.40 × to 12.31× with increase in *s* when *n* = 100 and 1000 respectively (see “*S2*.*1*.*2 n,s effect with fixed B*.*xlsx*” in Supplementary Data S2).

**Fig. 2.**
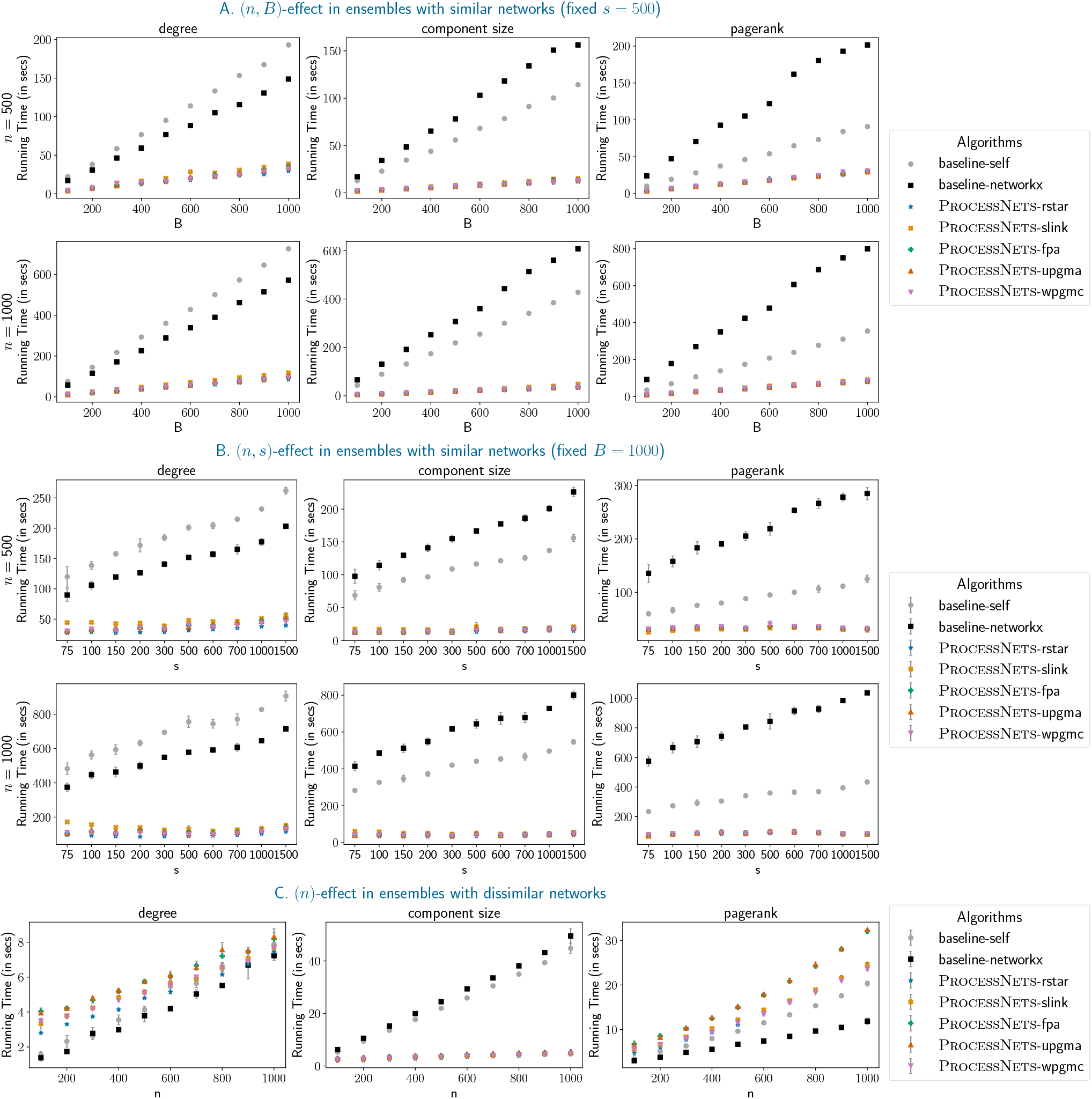
Results on Simulated Ensembles. A. Running times of baseline and our proposed framework to compute the network science measures of Simulated Ensemble I across varied *B*, when *n* is 500 or 1000 and sample size is fixed at *s* = 500. B. Similar results as in panel A for Simulated Ensemble I, but across varied *s*, when *n* is 500 or 1000, and bootstrap samples is fixed at *B* = 1000. This analysis was performed thrice, and the average running time across the three trials is shown, along with error bars indicating the standard deviation. C. Similar plots as in panel B, but for Simulated Ensemble II, across varied *n*. With the exception of a few cases, the error bars in panels B and C show minimal run-to-run variation in the running times; so we performed only one trial for the other analyses.

Among the ProcessNets-^*^, ProcessNets-rstar shows marginal improvements in the runtime at millisecond resolution for most of the scenarios; however, for computation of PR centrality, ProcessNets-slink also shows faster running times than the other configurations in several scenarios (see interactive plots and files in “*S2*.*1 Simulated Ensemble I*” in Supplementary Data S2).

### B. Results on Simulated Ensemble II (Comprising Dissimilar Networks)

As a negative control, we wanted to test the performance of our framework on ensembles with low similarity index. We varied *n* to generate multiple ensembles—each comprising networks generated independently using a degree sequence sampled from a power-law distribution. Given the low values of MPJS and smaller average edge count (i.e., in the range [126, 1391]) in these ensembles, the ProcessNets-^*^ perform poorly in computing degree and PR centrality (Fig. 2C and “*S2*.*2 Simulated Ensemble II*.*xlsx*” in Supplementary Data S2), as anticipated. An exception is observed for the component size computations, wherein the ProcessNets-^*^ runs significantly faster than the baseline-^*^, possibly due to the constant amortized time complexity of the incremental algorithm. Also, we note that for degree and PR centrality computations, `baseline-networkx` is the fastest.

### C. Real-World Ensembles on Muscle-Skeletal Dataset

Given the promising performance of our framework on simulated ensembles of similar networks, we wanted to inspect if the same also holds on real-world gene coexpression networks constructed from (boostrapped or subsampled) data on top-*n* varying genes of the GTEx Muscle Skeletal tissue (see Section II-F3).

For degree and component size computations, consistent with results in Simulated Ensemble I, the ProcessNets-^*^ outperform the baseline-^*^ across ensembles in all the three categories, except for a few scenarios when *n* = 100 (see Fig. 3). Mostly, the speedup increases with increase in *n* and *f*, as the running time of the ProcessNets-^*^ increase slowly as compared to that of the baseline-^*^. For example, for degree centrality, the speedup of ProcessNets-rstar over baseline-networkx increases with *n* from 0.53 × to 1.98 × for category ℬ *S*and 0.56 ×to 4.82 × for category *S*s. Also, the speedup for category *S sf* ranging between 2.72 × and 4.40 × is positively correlated to *f* (see files in “*S2*.*3*.*1 Muscle Skeletal*” in Supplementary Data S2).

**Fig. 3.**
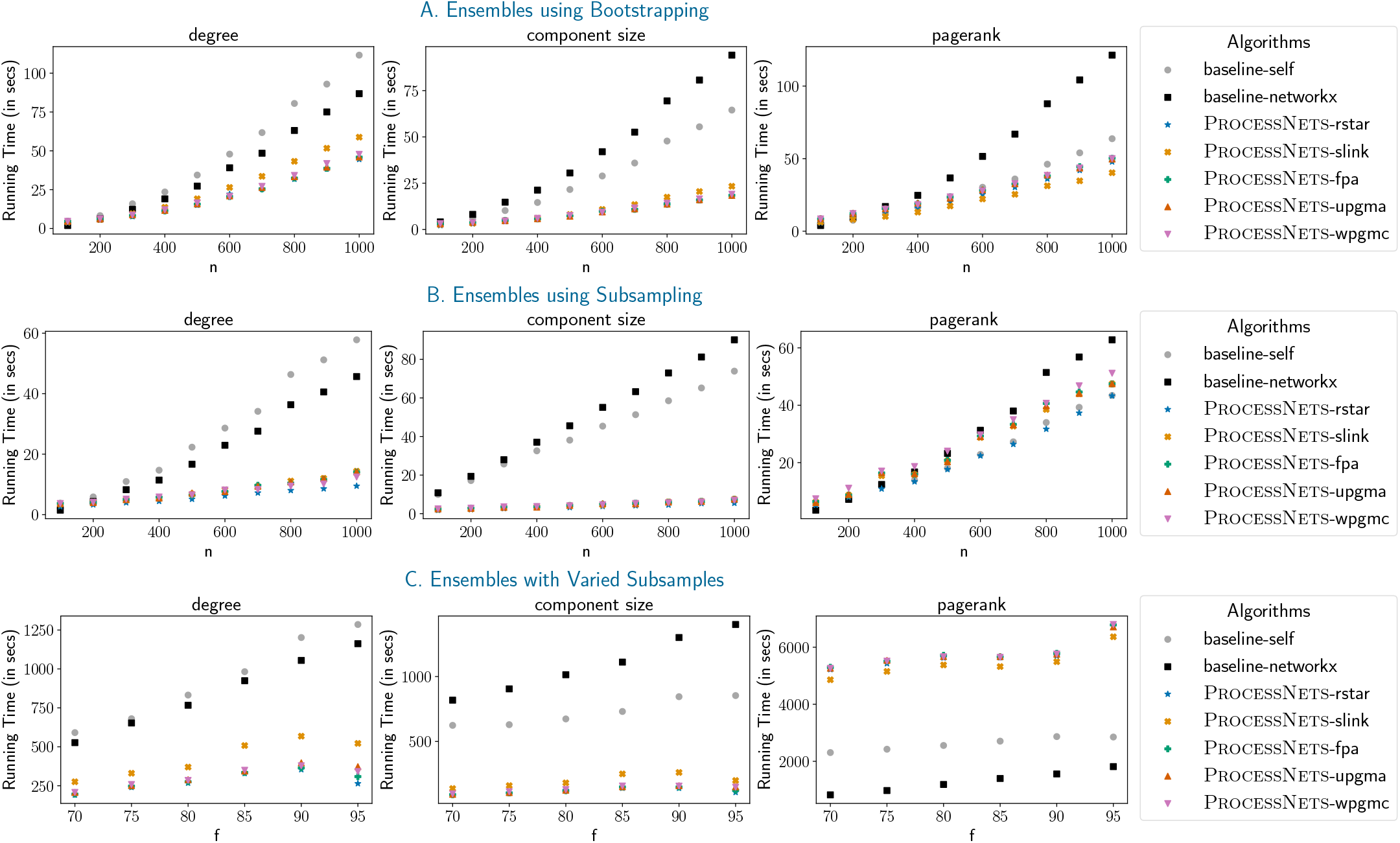
Results on Real-World Ensembles Constructed using GTEx Muscle-Skeletal Dataset. A. Plots depicting running times of different configurations to compute the network science measures across different number (*n*) of top-varying genes for ensembles in category ℬ*s*. B. Similar plots as in panel A. for ensembles in category *Ss* contructed using top-*n* varying genes. C. Similar plots as in panel A. for ensembles in category *Ssf*.

For PR centrality computations, we see interesting category-specific performance trends as follows:

- The results on category ℬ*s* ensembles, with average EC ranging in [352, 23272], are in accordance with Simulated Ensemble I results. ProcessNets-^*^ compute the central-ity measure faster than the baseline-^*^ and the speedup increases with *n* (Fig. 3A). For instance, the speedup of ProcessNets-slink over baseline-self increases from 0.65 × to 1.58 × with increase in *n* (“*S2*.*3*.*1A Category Bs*.*xlsx*” in Supplementary Data S2).
- Conversely, on category *Ss* ensembles (average EC [237, 12086]), ProcessNets-rstar and baseline-self execute faster than the other configurations except when *n* = 100 (Fig. 3B and “*S2*.*3*.*1B Category Ss*.*xlsx*” in Supplementary Data S2).
- On the other hand, the baseline-^*^ outperform the ProcessNets-^*^ on category *sf* ensembles with average EC in [103608, 258413] (Fig. 3C and “*S2*.*3*.*1C Category Ssf*.*xlsx*” in Supplementary Data S2).

Focusing on PR centrality, for ensembles where our framework did not outperform the baseline approach (e.g., category *Ss* or *Ssf* ensembles above), the low average edge count of these ensembles could be a likely reason (as detailed in Section IV-C2). However, our framework efficiently computes PR centrality for similar network ensembles with relatively large number of edges like category B*s* ensembles above.

### D. Real-World Ensembles on Several Tissues

We next wanted to examine if performance trends observed in the Muscle Skeletal tissue also holds for other tissues, focusing on scenarios where the network science measures are computed for all protein-coding genes filtered for the tissue (i.e., *n* ≈ 15, 000). Fig. 4 shows that the trends do hold across the 10 tested tissues, i.e., the ProcessNets-^*^ run faster than the baseline-^*^ for all tested scenarios (subject to the same exceptions involving PR centrality mentioned in the previous section). For instance, ProcessNets-rstar shows a speedup in the range [1.17, 1.72] × and [2.86, 3.97] × over baseline-networkx to compute degree centrality for ensembles in categories ℬ*s* and *Ss*, respectively (see files in “*S2*.*3*.*2 All Chosen Tissues*” in Supplementary Data S2). Additionally, results for PR centrality on category *Ss* ensembles of all tissues (obtained via 90% subsampling) are also consistent with results from previous section on category *Ssf* ensembles of the Muscle Skeletal tissue (restricted to *f* = 90% subsampling), as the number of nodes and average edge count of these ensembles are similar.

**Fig. 4.**
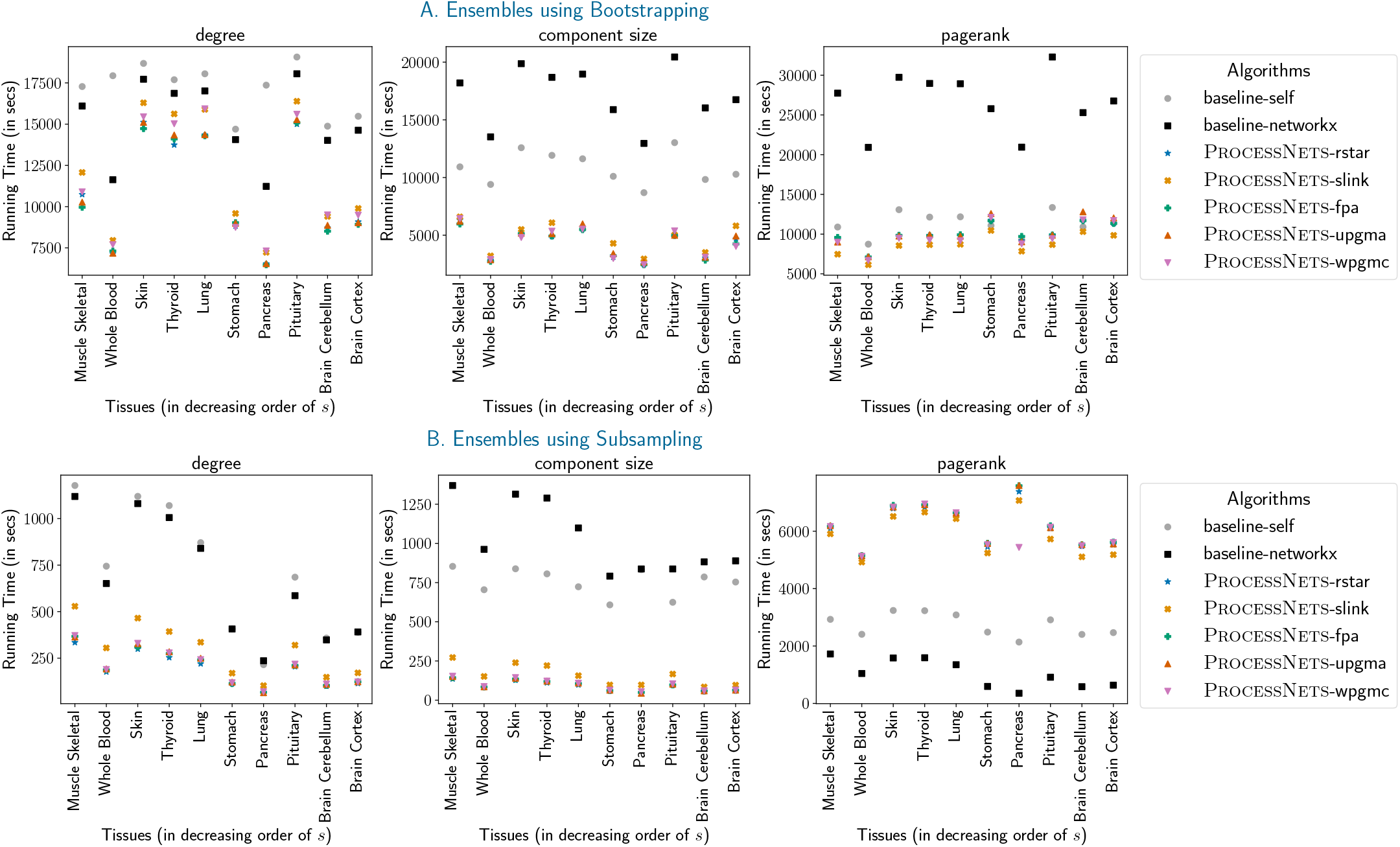
Results on Real-World Ensembles Constructed using Several GTEx Datasets. A. Plots depicting running times of different configurations to compute the network science measures across varied tissues for ensembles in category ℬ*s*. B. Similar plots as in panel A. for ensembles in category *Ss*.

## IV. Overall Inferences and Recommendations

### A. Effect of the nc-tree

Computation of the distance matrix in the hierarchical clustering based algorithms to construct the corresponding nc-trees requires pairwise analysis of the min-hash signatures among all networks in the ensemble (cf. Section II-C2), and is more time-consuming than constructing nc-trees using the rstar heuristic (Algorithm 1). For instance, while *T*_rstar_ for the category ℬ *s* and category *S s* ensembles on Whole Blood tissue is constructed in 2713 and 238 seconds respectively, construction of the other nc-trees require ≈ 7150 and ≈ 984 seconds respectively (see files in “*S2*.*3*.*2 All Chosen Tissues*” in Supplementary Data S2). However, since both rocessNets-rstar and ProcessNets-slink exhibit slightly better or comparable running times in computing the network measures, we recommend the use of rstar algorithm.

Our analysis on the breakdown of total running time (of certain real-world ensembles into the time for input, output and processing operations; see Appendix B.B and Table S2) provide definitive evidence that nc-trees, particularly *T*_rstar_, achieve a highly compact representation of all networks in an ensemble. This is especially true in ensembles with high MPJS, where compression factor as high as 8.1 × was observed. A highly compact representation leads to significant reduction in time for input operations. This speedup in input tasks coupled with the speedup in processing from using incremental algorithms contributes to the overall speedup of our framework.

### B. Effect of MPJS

ProcessNets-^*^ show higher running times relative to the baseline-^*^ in ensembles with highly dissimilar networks (MPJS ≈0.1), as evident from the results on Simulated nsemble II (Fig. 2C; degree and PR centrality). This result is seen possibly as very few edges are common among networks in these ensembles, hence new incremental edges are added along branches close to the leaves in their nc-trees; this results in a large number of incremental edges in the nc-trees. Based on these results, we recommend the use of baseline-^*^ in computation of network science measures in such scenarios.

### C. Effect of The Network Science Measure

To assess the effect of different network science measures on the efficiency of our framework, we now compare their performance trends on all ensembles of similar networks such as Simulated Ensemble I or real-world networks.

#### 3) Broad Benefits for Component Size and Degree Centrality

As previously noted, ProcessNets-^*^ show significantly faster execution time as compared to the baseline-^*^ to compute component size as well as degree centrality across different ensembles of similar networks. Specifically, for ensembles of similar networks with MPJS ≥ 0.2, ProcessNets-rstar computes component size and degree centrality the fastest among all seven configurations (Supplementary Data S3). This observation may be attributed to the structure of nc-trees. At any branching point *G* in *T*, if a connected component forms, then all edges in this connected component are not further processed during subsequent iterations of the incremental algorithm at branching points in the clade rooted at *G* (cf. Appendix A.C2). Conversely, the baseline-^*^ process this edge independently for every network in the tip of the same clade. This confers a significant advantage to the ProcessNets-^*^ over the baseline-^*^. A similar observation also holds for processing edges to compute degree centrality (cf. Appendix A.C1). Based on the empirical results and these interpretations, we recommend the use of ProcessNets-rstar to compute both connectivity- and degree-based measures in ensembles of similar networks.

#### 2) Conditional Benefit for PR Centrality

For computing PR centrality on networks over a large number of genes (i.e., *n* ≈ 15, 000), ProcessNets-^*^ runs faster than baseline-^*^ for certain real-world ensembles (e.g., Category ℬ*s*), and slower for certain other ensembles (e.g., Category *Ss*). To dissect this trend further, we explored whether MPJS or average edge count of the ensemble could be contributing factors. The effect of MPJS mentioned in Section IV-B above is not aligned with this trend, however average edge count was highly correlated to the speedup achieved by ProcessNets-^*^ (Fig. 5). Specifically, for ensembles with at least 3 × 10^6^ average edge count, the speedup of ProcessNets-slink was at least 1.39× over the fastest baseline-^*^. If the average edge count is lower than this threshold, we hypothesize that the overheads associated with ProcessNets-^*^ is not justifiable relative to computing PR using baseline-^*^. To test this hypothesis further, we perturbed the edge count of the two real-world Muscle Skeletal ensembles (i.e., changed Category *Bs* ensemble from default high average edge count to a low count by varying FDR, and vice versa for Category *S s* ensemble (see Table S3)), and found that the hypothesis indeed holds (see Appendix B.C and Fig. S2). Based on these inferences and other results on PR centrality, we recommend ProcessNets-slink for ensembles of networks with a large number of nodes and edges, ProcessNets-rstar for network ensembles with a small number of nodes, and baseline-networkx for other scenarios.

**Fig. 5.**
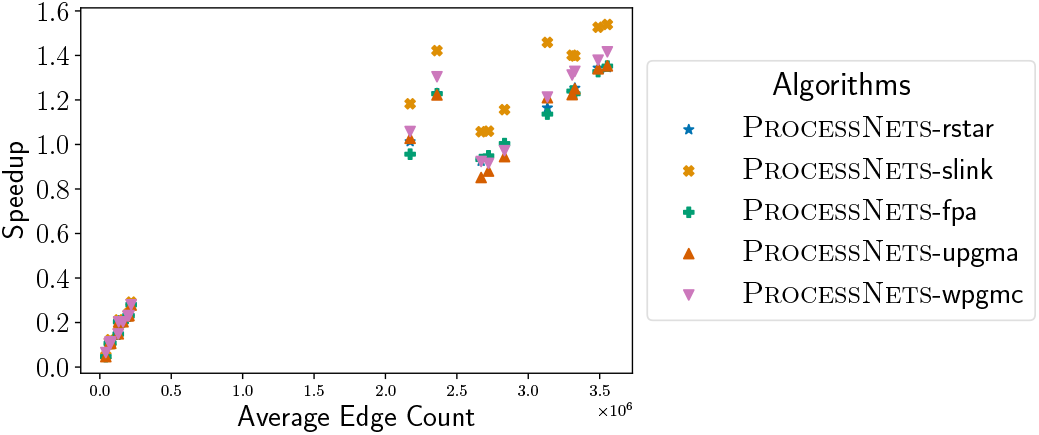
Speedup achieved by different ProcessNets-^*^ relative to the fastest baseline-^*^ in computing PR centrality with respect to average edge count in real-world ensembles constructed on GTEx tissues used across all analyses. Note that only real-world ensembles constructed using all filtered protein-coding genes (i.e., ensembles used in Fig. 4 and Fig. S2B) are used to generate this scatter plot.

## V. Conclusion and Future Works

In this study, we address the computational bottleneck of integrating information (graph/node-level properties) from numerous networks pertaining to a biological system. These networks are often similar to each other, as they are all related to the same biological system and inferred from an underlying set of coherently related datasets. We propose the ProcessNets framework to harness this similarity among networks in an ensemble by representing them as a compact DS nc-tree and running incremental graph algorithms along the edges of this nc-tree. Systematic analytical and empirical evaluations of the efficiency of our framework relative to a baseline approach across varied network science measures and simulated/real-world ensembles led to the following key findings and recommendations:

- ProcessNets-^*^ configurations compute network science measures (such as degree centrality, component size, and PR centrality) faster than the baseline-^*^ in ensembles with high MPJS or average edge count. As a result, we can now compute these measures for a larger number of input networks under the same budget of compute resources (such as the number of parallel cores).
- ProcessNets-rstar shows faster running times than other configurations for computing degree- and connectivity-based measures. Similarly, the ProcessNets-slink con-figuration shows higher speedup in computing PR centrality in ensembles comprising networks with a large number of nodes and edges.
- Our compact and reusable DS, nc-tree, enables efficient storage, reconstruction, and analysis of an input ensemble, as long as the ensembles comprise similar networks. The compression factor achieved by nc-tree in storing all networks in an ensemble results in not only less disk space to store the ensemble, but also significantly less time for input operations (compared to reading the networks one at a time).

To the best of our knowledge, ProcessNets is the first algorithmic framework to harness the similarity among a set of biological networks to speedup property computations on these graphs. Our ProcessNets framework utilizes a phylogeny-like DS called nc-tree to achieve this (space and time) speedup. Other bioinformatics studies have utilized phylogeny-like structures for compaction, but for compressing the storage of other objects such as different suffixes of a string (linear-space suffix trees [22], [23]), and all-vs-all pairwise distances among a large set of microbial genomes (subquadratic-space PLATO [24]). In our GTEx applications, we have used our framework to speedup bootstrap analysis of network properties. Other bioinformatics studies have used efficient algorithms to speedup bootstrap analysis, but they have focused on other problems such as quantifying the expression of gene isoforms using equivalence-class-based grouping of RNAseq reads (Kallisto [25]).

Despite the promising speedups achieved with our framework, there are some limitations of our study. While this study focuses on our framework’s efficiency primarily on three specific network science measures, its performance on other computationally expensive measures is yet to be explored. Nevertheless, we believe the ProcessNets framework has the potential to efficiently compute such measures with properly chosen batch incremental dynamic graph algorithms that are based on compact and information-rich intermediate DSs. Another limitation of our framework is its overhead on integrating dissimilar networks or sparse networks with very few nodes and edges—in these scenarios, we recommend using the baseline over our approach. A final caveat pertains to comparisons of the different configurations done under a sequential implementation mode (see Appendix A.E); specifically, a parallel implementation of our framework, say by executing a forest of nc-trees simultaneously, has not been compared with an embarrassingly parallel implementation of the baseline approach. However, if the number of nc-trees in the forest matches the number of available parallel cores, we believe that the speedup of our framework over baseline reported in this study should largely hold for the parallel implementation mode also. This belief relies on whether the different trees in the forest can be optimized to achieve balance, in terms of the number of networks in the ensemble that each tree represents. Addressing this and related optimization problems remain an important direction for future study.

All empirical analyses in this study were performed on ensembles comprising transcriptomic networks. However, our framework is readily applicable to ensembles of other omics-based biomolecular networks, and even non-molecular networks such as ecological [26], [27], species interaction [28], [29], or brain networks [30]–[33] where network science measures are routinely calculated, with our framework’s speedup dependent on the similarity of the input networks. We antici-pate that our ProcessNets framework based on nc-trees and incremental algorithms can serve as a foundation for future methodological advances in scalable analysis of large-scale biological network ensembles.

## Code AND Data availability

The code for the ProcessNets framework and all other associated analyses is available at https://github.com/BIRDSgroup/ProcesssNets. Running times of several configurations across varied settings is available at https://drive.google.com/drive/folders/1GJ6Nj2qO_Cyy5uP_dWZr3oGaqW-OW_tE?usp=sharing.

## Acknowledgments

The authors thank Saish Jaiswal for reviewing the manuscript. The authors also acknowledge the assistance provided by Shaun Mathew, Shankar Balajee, T. M. Kumaresan, Ayon Ghosh, Snehadeep Gayen, Ayman Akhter, Dasari Himakar Sai, Guntupalli Ashish Sai, T Sai Krishna, Gandikota Sai Pradhyumna, Pranav B, Thiruvarul P, Gokulakrishnan R, Ganesh T and Arjun Vikas Ramesh in implementing existing incremental algorithms for closeness centrality. AI models such as ChatGPT and Gemini were utilized for copy-editing purposes (including polishing certain phrases or paragraphs) and minimal assistance with coding certain functions, and the Undermind AI platform was used to assist the literature review process. All AI-generated content were verified manually before inclusion in this work.

## Author contributions

This work is performed as part of the doctoral thesis of SM with extensive inputs from MN.

## References

[1] V. Gligorijević and N. Pržulj, “Methods for biological data integration: perspectives and challenges,” Journal of The Royal Society Interface, vol. 12, no. 112, 2015.

[2] K. Mitra, A.-R. Carvunis, S. K. Ramesh, and T. Ideker, “Integrative approaches for finding modular structure in biological networks,” Nature Reviews Genetics, vol. 14, no. 10, pp. 719–732, 2013.

[3] M. Picard, M.-P. Scott-Boyer, A. Bodein, O. Périn, and A. Droit, “Integration strategies of multi-omics data for machine learning analysis,” Computational and Structural Biotechnology Journal, vol. 19, pp. 3735–3746, 2021.

[4] G. Didier, C. Brun, and A. Baudot, “Identifying communities from multiplex biological networks,” PeerJ, vol. 3, p. e1525, 2015.

[5] The GTEx Consortium, “The GTEx Consortium atlas of genetic regulatory effects across human tissues,” Science, vol. 369, no. 6509, pp. 1318–1330, 2020.

[6] Cancer Genome Atlas Research Network, J. N. Weinstein, E. A. Collisson, G. B. Mills, K. R. M. Shaw, B. A. Ozenberger, K. Ellrott Shmulevich, C. Sander, and J. M. Stuart, “The Cancer Genome Atlas Pan-Cancer analysis project,” Nature Genetics, vol. 45, no. 10, pp. 1113–1120, 2013.

[7] B. B. Sun, J. Chiou, M. Traylor, C. Benner, Y.-H. Hsu, T. G. Richardson, P. Surendran, A. Mahajan, C. Robins, S. G. Vasquez-Grinnell, L. Hou, E. M. Kvikstad, O. S. Burren, J. Davitte, K. L. Ferber, C. E. Gillies, Å. K. Hedman, S. Hu, T. Lin, R. Mikkilineni, R. K. Pendergrass, C. Pickering, B. Prins, D. Baird, C.-Y. Chen, L. D. Ward, A. M. Deaton, S. Welsh, C. M. Willis, N. Lehner, M. Arnold, M. A. Wörheide, K. Suhre, G. Kastenmüller, A. Sethi, M. Cule, A. Raj, Alnylam Human Genetics, AstraZeneca Genomics Initiative, Biogen Biobank Team, Bristol Myers Squibb, Genentech Human Genetics, GlaxoSmithKline Genomic Sciences, Pfizer Integrative Biology, Population Analytics of Janssen Data Sciences, Regeneron Genetics Center, L. Burkitt-Gray, E. Melamud, M. H. Black, E. B. Fauman, J. M. M. Howson, H. M. Kang, M. I. McCarthy, P. Nioi, S. Petrovski, R. A. Scott, E. N. Smith, S. Szalma, D. M. Waterworth, L. J. Mitnaul, J. D. Szustakowski, B. W. Gibson, M. R. Miller, and C. D. Whelan, “Plasma proteomic associations with genetics and health in the UK Biobank,” Nature, vol. 622, no. 7982, pp. 329–338, 2023.

[8] M. Newman, Networks: An Introduction, 2nd ed. Oxford University Press, 2018.

[9] S. Gómez, “Centrality in Networks: Finding the Most Important Nodes,” in Business and Consumer Analytics: New Ideas. Springer, 2019, pp. 401–433.

[10] M. Wang, H. Wang, and H. Zheng, “A Mini Review of Node Centrality Metrics in Biological Networks,” International Journal of Network Dynamics and Intelligence, vol. 1, no. 1, pp. 99–110, 2022.

[11] Z. Chen, K. Liang, L. Yuan, W. Zhang, and Z. Yang, “Recent Advances in Efficient Dynamic Graph Processing,” Applied Sciences, vol. 15, no. 11, p. 6003, 2025.

[12] J. H. Camin and R. R. Sokal, “A Method for Deducing Branching Sequences in Phylogeny,” Evolution, vol. 19, no. 3, pp. 311–326, 1965.

[13] W. H. Day, D. S. Johnson, and D. Sankoff, “The computational complexity of inferring rooted phylogenies by parsimony,” Mathematical Biosciences, vol. 81, no. 1, pp. 33–42, 1986.

[14] S. Mahapatra, M. Narayanan, and N. S. Narayanaswamy, “Parameterized algorithms for the Steiner arborescence problem on a hypercube,” Acta Informatica, vol. 62, no. 1, p. 6, 2025.

[15] B. S. Everitt, S. Landau, M. Leese, and D. Stahl, Cluster Analysis, 5th ed. Wiley Series in Probability and Statistics, 2011.

[16] J. Leskovec, A. Rajaraman, and J. D. Ullman, Mining of Massive Datasets, 3rd ed. Cambridge University Press, 2020.

[17] T. H. Cormen, C. E. Leiserson, R. L. Rivest, and C. Stein, Introduction to Algorithms, 3rd ed. The MIT Press, 2009.

[18] C. C. Paige and M. A. Saunders, “Solution of Sparse Indefinite Systems of Linear Equations,” SIAM Journal on Numerical Analysis, vol. 12, no. 4, pp. 617–629, 1975.

[19] L. C. Freeman, “Centrality in social networks conceptual clarification,” Social Networks, vol. 1, no. 3, pp. 215–239, 1978.

[20] A. E. Sariyüce, K. Kaya, E. Saule, and Ü. V. Ç atalyiirek, “Incremental algorithms for closeness centrality,” in 2013 IEEE International Conference on Big Data. IEEE, 2013, pp. 487–492.

[21] S. Mahapatra, N. A. Subramanian, and M. Narayanan, “Uncertainty-Aware Gene Rankings Reveal Key Players in Coexpression Networks,” bioRxiv, 2025.

[22] P. Compeau and P. Pevzner, Bioinformatics Algorithms: An Active Learning Approach, 3rd ed. Active Learning Publishers, 2020.

[23] D. Gusfield, Algorithms on Strings, Trees, and Sequences: Computer Science and Computational Biology, 1st ed. Cambridge University Press, 1997.

[24] L. Ackermann, P. Peterlongo, and K. Břinda, “Subquadradic storage of pairwise distance matrices of large microbial genome collections.” RECOMB 2026 Best Poster Award, 2026.

[25] N. L. Bray, H. Pimentel, P. Melsted, and L. Pachter, “Near-optimal probabilistic RNA-seq quantification,” Nature Biotechnology, vol. 34, no. 5, pp. 525–527, 2016.

[26] D. R. Hemprich-Bennett, H. F. M. Oliveira, S. C. Le Comber, S. J. Rossiter, and E. L. Clare, “Assessing the impact of taxon resolution on network structure,” Ecology, vol. 102, no. 3, p. e03256, 2021.

[27] M. J. Michalska-Smith and S. Allesina, “Telling ecological networks apart by their structure: A computational challenge,” PLOS Computational Biology, vol. 15, no. 6, p. e1007076, 2019.

[28] S. H. Fard and E. Dolson, “The Robustness of Structural Features in Species Interaction Networks,” 2025.

[29] C. Brimacombe, K. Bodner, M. Michalska-Smith, T. Poisot, and M.-J. Fortin, “Shortcomings of reusing species interaction networks created by different sets of researchers,” PLOS Biology, vol. 21, no. 4, p. e3002068, 2023.

[30] F. Xu, S. Garai, D. Duong-Tran, A. J. Saykin, Y. Zhao, and L. Shen, “Consistency of Graph Theoretical Measurements of Alzheimer’s Disease Fiber Density Connectomes Across Multiple Parcellation Scales,” in 2022 IEEE International Conference on Bioinformatics and Biomedicine (BIBM). IEEE, 2022, pp. 1323–1328.

[31] T. Welton, C. S. Constantinescu, D. P. Auer, and R. A. Dineen, “Graph Theoretic Analysis of Brain Connectomics in Multiple Sclerosis: Reliability and Relationship with Cognition,” Brain Connectivity, vol. 10, no. 2, pp. 95–104, 2020.

[32] S. Hanalioglu, S. Bahadir, I. Isikay, P. Celtikci, E. Celtikci, F.-C. Yeh, K. K. Oguz, and T. Khaniyev, “Group-Level Ranking-Based Hubness Analysis of Human Brain Connectome Reveals Significant Interhemispheric Asymmetry and Intraparcel Heterogeneities,” Frontiers in Neuroscience, vol. 15, p. 782995, 2021.

[33] P. Borrelli, C. Cavaliere, M. Salvatore, J. Jovicich, and M. Aiello, “Structural Brain Network Reproducibility: Influence of Different Diffusion Acquisition and Tractography Reconstruction Schemes on Graph Metrics,” Brain Connectivity, vol. 12, no. 8, pp. 754–767, 2022.

