## Supplementary Information for "ProcessNets: Towards an efficient approach for ensemble analysis of biological networks"

This work has been submitted to the IEEE for possible publication. Copyright may be transferred without notice, after which this version may no longer be accessible.

### ProcessNets: Towards an efficient approach for ensemble analysis of biological networks (Supplementary Information)

Sugyani Mahapatra, and Manikandan Narayanan

A.C1 Algorithms for Degree Centrality 2
A.C2 Algorithms for Component Size 2
A.C3 Algorithms for PR Centrality 2
A.C4 Algorithms for Closeness Centrality 3
A.D Construction of Coexpression Network from Gene Expression Dataset . . . . . 3
A.E Implementation Details of Ensemble Construction, and ProcessNets-* and baseline-* Configurations . . . . . 4

**Appendix B: Supplementary Results** 5
B.A Computation of Closeness Centrality . . 5
B.B Dissecting ProcessNets Speedup using I/O (Input/Output) vs. Processing Times . . . . . 5
B.C Dependence of Running Time to Compute PR Centrality on Average Edge Count 5

**Appendix C: Supplementary Data** 7

**References** 7

The authors are with Department of Computer Science & Engineering, Centre for Integrative Biology and Systems medicinE, and Wadhwani School of Data Science & AI, Indian Institute of Technology (IIT) Madras, Chennai, India. Second author is also affiliated to Sudha Gopalakrishnan Brain Centre, IIT Madras, Chennai, India.

### Appendix A
### Supplementary Methods

#### A. Pseudocode of Our ProcessNets Framework

The pseudocode of our proposed ProcessNets framework, which proceeds in three phases to compute a network science measure $\mathcal{P}$ for each network in the input ensemble $\mathcal{G}$ (as explained in Section II-B2 in the main text), is shown in Algorithm S1.

**Algorithm S1** The ProcessNets Algorithm

**Input:** An ensemble of networks $\mathcal{G} = \{G_1 = (V, E_1), G_2 = (V, E_2), \ldots, G_B = (V, E_B)\}$
**Output:** The network science measure $\mathcal{P}$ in each of the networks, i.e., $\mathcal{P}(G_1), \mathcal{P}(G_2), \ldots, \mathcal{P}(G_B)$

```
▷ Phase P1: NC-tree Construction
T ← A_tree(G)

▷ Phase P2: Measurements of the Root Graph P(G_{1,...,B})
P(G_{1,...,B}) ← A_static(G_{1,...,B})

▷ Phase P3: Hierarchical Computation at Branching Points. Although
  any top-down tree traversal algorithm can be used, we adopt a level-
  order traversal for space efficiency.
Q ← EmptyQueue()
Enqueue(Q, G_{1,...,B})
while Q ≠ ∅ do
    G_parent ← Dequeue(Q)
    for all child G_child of G_parent do
        P(G_child) ← A_dynamic(P(G_parent), E_child \ E_parent)
        Enqueue(Q, G_child)
    end for
    [optional] Delete P(G_parent)
end while
```

#### B. Distance Metric Computation using Minhash Signatures

Let $\mathcal{G}$ be the input ensemble of networks, each defined on a set of $n$ nodes. We generated 10 random permutations $(\pi_1, \pi_2, \ldots, \pi_{10})$ of the set comprising all possible $\binom{n}{2}$-edges on these nodes. For every network $G = (V, E) \in \mathcal{G}$ and each permutation $\pi_i$, we define the $i^{\text{th}}$ signature of $G$ as the first edge in $\pi_i$ that is also present in $E$. For each network $G \in \mathcal{G}$, this process resulted in a minhash signature [1] of length 10 (represented as a vector $\in \left\{1, 2, \ldots, \binom{n}{2}\right\}^{10}$). Distance between a pair of networks $G_i$ and $G_j$ is defined as the Hamming distance between their corresponding minhash signatures. Note that for a position $i \in \{1, 2, \ldots, 10\}$, the probability that $G_i$ and $G_j$ agree on the $i$th position of their minhash vectors is simply the Jaccard similarity index between their edge sets (i.e., $\frac{|E_i \cap E_j|}{|E_i \cup E_j|}$).

#### C. Algorithms and Implementations of Network Science Measures

In this section, we describe the different algorithms used to compute four network science measures in our empirical analysis and their implementation details.

*1) Algorithms for Degree Centrality:* We used the straightforward approach, wherein we iterate over every edge and increment the degree of the endpoints by 1, as the static algorithm to compute degree centrality for the `baseline`-self and ProcessNets-*. The resulting degree values are stored in an array indexed by node identifiers; each node is assigned a unique integer index, and a lookup table maintains the mapping between indices and node names. At every branch of the NC-tree, we iteratively update the array containing the degrees of the parent graph to compute the degrees of the child graph, i.e., for every new incremental edge, the corresponding entries of the endpoints in the array are incremented by 1. This incremental algorithm is used for all the five ProcessNets-*. For the `baseline`-networkx configuration, the NetworkX graph is constructed for each network and the `degree` function is used.

*2) Algorithms for Component Size:* In the `baseline-*`, we iteratively apply depth-first search (DFS) (as recommended in [2]) on an unvisited node $u$ to generate the set of nodes in the component containing $u$; the component size of every node in this component is the size of this set. While the function to apply the DFS was implemented in the `baseline`-self configuration, `dfs_preorder_nodes` function was used on the constructed NetworkX graph in the `baseline`-networkx configuration.

For the ProcessNets-*, we used the union-find data structure (union-by-size) with path compression [2] (both for static and incremental algorithms) to compute the component sizes. It may be noted here that although the algorithm inspects every edge sequentially, the union operation is performed only for a few edges (connecting disjoint components) and the measurements are recomputed only if such an operation has been performed. This ensures that once the components are connected at any branching point of the NC-tree, no further operations are performed; we call this as partial batch analysis. Similar to degree centrality, we use an array indexed by node identifiers to store the component sizes here as well.

*3) Algorithms for PR Centrality:* To optimize the intermediate DS required at each branching point, we adapted the incremental algorithm for PR centrality in [3] following an approach used in the incremental algorithm for Katz centrality in [4] by performing a change of variables.

Let $\widetilde{D}_I \in \mathbb{R}^{n \times n}$ denote the diagonal matrix with $\widetilde{D}_I[u, u] = \max\{\deg_I[u], 1\}$; here $\deg_I[u]$ is the degree (out-degree for directed graphs) of the node $u$ in the graph $G_I$. Then, the PR centrality of all nodes in $G_I$, represented by the column-vector $\vec{x}_I$ (indexed by node identifiers similar to that in degree centrality), is computed as the solution to the linear system $(I_n - \alpha A_I \widetilde{D}_I^{-1})\vec{x}_I = \frac{(1-\alpha)}{n}\vec{1}$; here $A_I$ and $I_n$ are the adjacency matrix of $G_I$ and the identity matrix with $n$ dimensions respectively. $\alpha$ represents the damping factor (usually set to 0.85). Note that, for this algorithm only, the $(i, j)^{\text{th}}$ entry of $A_I$ equals 1 only if there is an edge from $j$ to $i$ in directed graphs, as per the notation followed in [5].

However, since our ensembles consist of large-scale networks with many nodes, we use a slightly modified (unnormalized) form of the original linear system, given by $(I_n - \alpha A_I \widetilde{D}_I^{-1})\vec{x}_I = (1 - \alpha)\vec{1}$ in all configurations except `baseline`-networkx. It may be noted here that without the $1/n$ normalization factor, the teleportation vector $\frac{1}{n}\vec{1}$ is no longer a probability vector, implying that the iteration does not correspond to a Markov chain and the usual random walk interpretation is lost. Nevertheless, since the matrix $(I_n - \alpha A_I \widetilde{D}_I^{-1})$ is still invertible, this unnormalized form of the linear system still has a unique solution and power iterations converge to it; the resulting vector is simply a scaled version (by a factor of $n$) of the vector containing the actual PR values.

Our adapted algorithm replaces the vector $\vec{x}_I$ with $\vec{y}_I$, defined as $\vec{y}_I = \widetilde{D}_I^{-1}\vec{x}_I$, and computes $\vec{y}_I$ by solving the linear system $M_I\vec{y}_I = (1-\alpha)\vec{1}$, where $M_I = \widetilde{D}_I - \alpha A_I$, using the MINRES iterative solver [6] with tolerance $10^{-5}$. It may be noted here that we use compressed sparse row (CSR) matrix representation to store the matrices and the `minres` function for sparse matrices provided by `scipy.sparse` package to solve this linear system of equations. Finally, $\vec{x}_I = \widetilde{D}_I\vec{y}_I$ can be simply computed as the dot product between the diagonal of $\widetilde{D}_I$ and $\vec{y}_I$ i.e., $\vec{x}_I[u] = \max\{\deg_I[u], 1\}\ \vec{y}_I[u]$. We use this algorithm as our static algorithm in all configurations except `baseline`-networkx.

For the incremental algorithm, we store the matrix $M_I$, and the vectors $\deg_I$ and $\vec{y}_I$ as intermediate DS at every branching point $G_I$ of the NC-tree. Suppose the edge set $E_\Delta$ is inserted to $G_I$ to obtain $G_J$ along the branch $e = (G_I, G_J)$, and suppose $\vec{y}_J = \vec{y}_I - \vec{y}_\Delta$; here the vector $\vec{y}_\Delta$ stores the updates required to the actual centrality values and we aim to compute this vector with faster convergence. Similarly, we also break the matrix $M_J$ into two components as $M_J = M_I + M_\Delta$. Since $M_I\vec{y}_I = (1-\alpha)\vec{1}$ (we assume exact solutions without residuals due to the very small tolerance), $\vec{y}_\Delta$ can be computed as follows:

$$
\begin{aligned}
M_J\vec{y}_J = (M_I + M_\Delta)(\vec{y}_I - \vec{y}_\Delta) &= (1-\alpha)\vec{1} \\
\implies M_I\vec{y}_I + M_\Delta\vec{y}_I - (M_I + M_\Delta)\vec{y}_\Delta &= (1-\alpha)\vec{1} \\
\implies M_\Delta\vec{y}_I - M_J\vec{y}_\Delta &= 0 \\
\implies M_J\vec{y}_\Delta = M_\Delta\vec{y}_I &
\end{aligned}
$$

Now, we use the notation $A_\Delta$ to represent the adjacency matrix of the network constructed on the edge set $E_\Delta$ and define $\widetilde{D}_\Delta$ as $\widetilde{D}_\Delta = \widetilde{D}_J - \widetilde{D}_I$ to compute $M_\Delta$ as:

$$M_\Delta = M_J - M_I = (\widetilde{D}_J - \alpha A_J) - (\widetilde{D}_I - \alpha A_I) = \widetilde{D}_\Delta - \alpha A_\Delta$$

The pseudocode of this incremental algorithm has been summarized below in Algorithm S2; the above description proves the correctness of both the static and incremental algorithm.

Note that although the worst-case time complexity is $\mathcal{O}(n^3)$, in practice our proposed incremental algorithm runs faster especially in similar network ensembles comprised of dense networks, possibly due to the CSR matrix representations. Our proposed incremental algorithm also provides the flexibility to initialize $\vec{y}_\Delta$ with either $\vec{0}$ or $\vec{y}_I$ depending on the number of non-zero elements in $M_\Delta$ ensuring faster convergence as compared to the static algorithm.

Finally, for the `baseline`-networkx configuration, we use the `pagerank` function on the constructed NetworkX graphs; it may be noted here that, unlike our other configurations, this function treats all input networks as directed and returns normalized centrality values.

**Algorithm S2** Incremental PageRank centrality algorithm applied to the branch $e = (G_I, G_J)$ of the NC-tree

**Input:** Intermediate DS of $G_I$: $M_I$, $\deg_I$ and $\vec{y}_I$, New edges inserted to $G_I$ to obtain $G_J$: $E_\Delta$
**Output:** If $G_J$ is a tip, then $\vec{x}_J$ else $y_J$

```
Convert E_Δ into an adjacency matrix A_Δ and compute deg_Δ.
deg_J ← deg_I + deg_Δ.
Compute D̃_I and D̃_J from deg_I and deg_J respectively.
D̃_Δ ← D̃_J − D̃_I.
M_Δ ← D̃_Δ − αA_Δ.
M_J ← M_I + M_Δ.
Solve the linear equation M_J y_Δ = M_Δ y_I using the MINRES iterative solver [6] with tolerance 10^−5. The initial value for y_Δ may be set to zero or y_I depending on the dataset.
y_J ← y_I − y_Δ
if G_J is a tip then
    return deg_J · y_J
else
    return y_J
end if
```

*4) Algorithms for Closeness Centrality:* We first note that this centrality measure was computed only with `baseline`-networkx and the PROCESSNETS-*. The `closeness_centrality` (uses Freeman's Algorithm from [7]) and `incremental_closeness_centrality` (uses the Work-filtering algorithm from [8]) functions from the NetworkX package were used as the static and incremental algorithms respectively to compute the measure. However, this incremental algorithm depends on the internal structure of the original graph instead of intermediate DS and does not process the incremental edges as a batch. As expected, performance of our framework was subpar for this measure even on smaller simulated networks, and hence detailed analyses on the same was not done (see Appendix B.A for all analysis on closeness centrality).

##### D. Construction of Coexpression Network from Gene Expression Dataset

Traditional tools/packages like WGCNA [9], GWENA [10], CoGTEx [11] and GeneFriends [12] can be used for construction of the coexpression network on a gene expression dataset. However, WGCNA and GWENA assume a power-law degree distribution among the genes, GeneFriends depends on a binomial model to compute the probability of co-occurrence, and CoGTEX uses aggregate correlation across tissues and datasets to compute a coexpression network. To avoid such assumptions, we used a simple network construction process as in [13]. Spearman correlation coefficient and the related $p$-value between every pair of genes (nodes in the network) was calculated, following which the $p$-values were adjusted using Benjamini-Hochberg FDR correction. Finally, edges were added between pairs of genes that satisfied the 1% FDR cutoff on the adjusted $p$-values.

TABLE S1
RUNNING TIME COMPARISON BETWEEN MULTI-CORE VS. SINGLE-CORE BENCHMARKING ON TWO ENSEMBLES, EACH COMPRISED OF $B = 1000$ NETWORKS. THE REPORTED VALUES ARE RUNNING TIMES IN MINUTES:SECONDS FORMAT, AND THE FASTEST RUNNING TIME IN EACH COLUMN IS SHOWN IN BOLD-FACED FONT.

(a) Degree Centrality

| Datasets → Configurations ↓ | Simulated Ensemble I ($n = 800$, $s = 700$) | | Simulated Ensemble II ($n = 800$) | |
|---|---|---|---|---|
| | Multi-core | Single-core | Multi-core | Single-core |
| | MPJS $= 0.76$ Avg. EC. $= 92,761.33$ | MPJS $= 0.76$ Avg. EC. $= 91,430.77$ | MPJS $= 0.10$ Avg. EC. $= 1,109.62$ | MPJS $= 0.10$ Avg. EC. $= 1,106.26$ |
| baseline-self | 8:12.309 | 7:18.176 | 0:06.510 | 0:06.087 |
| baseline-networkx | 6:29.338 | 5:35.948 | **0:05.439** | **0:05.559** |
| ProcessNets-rstar | **1:11.682** | **1:00.024** | 0:06.076 | 0:05.588 |
| ProcessNets-slink | 1:32.113 | 1:16.262 | 0:06.410 | 0:05.658 |
| ProcessNets-fpa | 1:19.522 | 1:07.730 | 0:07.153 | 0:06.252 |
| ProcessNets-upgma | 1:20.851 | 1:09.319 | 0:07.157 | 0:06.272 |
| ProcessNets-wpgmc | 1:16.159 | 1:04.311 | 0:06.436 | 0:05.629 |

(b) Component Size

| Datasets → Configurations ↓ | Simulated Ensemble I ($n = 800$, $s = 700$) | | Simulated Ensemble II ($n = 800$) | |
|---|---|---|---|---|
| | Multi-core | Single-core | Multi-core | Single-core |
| | MPJS $= 0.76$ Avg. EC. $= 92,761.33$ | MPJS $= 0.76$ Avg. EC. $= 91,430.77$ | MPJS $= 0.10$ Avg. EC. $= 1,109.62$ | MPJS $= 0.10$ Avg. EC. $= 1,106.26$ |
| baseline-self | 4:54.259 | 4:16.206 | 0:34.980 | 0:34.412 |
| baseline-networkx | 7:16.502 | 6:05.157 | 0:38.077 | 0:38.248 |
| ProcessNets-rstar | **0:25.390** | **0:22.321** | **0:03.848** | **0:03.478** |
| ProcessNets-slink | 0:33.989 | 0:27.777 | 0:04.181 | 0:03.818 |
| ProcessNets-fpa | 0:29.266 | 0:25.474 | 0:04.948 | 0:04.257 |
| ProcessNets-upgma | 0:29.569 | 0:26.049 | 0:04.795 | 0:04.268 |
| ProcessNets-wpgmc | 0:27.824 | 0:23.524 | 0:04.186 | 0:03.711 |

(c) PR Centrality

| Datasets → Configurations ↓ | Simulated Ensemble I ($n = 800$, $s = 700$) | | Simulated Ensemble II ($n = 800$) | |
|---|---|---|---|---|
| | Multi-core | Single-core | Multi-core | Single-core |
| | MPJS $= 0.76$ Avg. EC. $= 92,761.33$ | MPJS $= 0.76$ Avg. EC. $= 91,430.77$ | MPJS $= 0.10$ Avg. EC. $= 1,109.62$ | MPJS $= 0.10$ Avg. EC. $= 1,106.26$ |
| baseline-self | 3:54.490 | 3:23.469 | 0:15.552 | 0:14.986 |
| baseline-networkx | 10:02.111 | 8:27.490 | **0:10.112** | **0:09.359** |
| ProcessNets-rstar | **0:58.017** | **0:44.333** | 0:18.366 | 0:15.820 |
| ProcessNets-slink | 1:01.234 | 0:50.969 | 0:18.849 | 0:16.038 |
| ProcessNets-fpa | 1:00.786 | 0:48.430 | 0:23.982 | 0:20.482 |
| ProcessNets-upgma | 1:01.415 | 0:48.075 | 0:24.377 | 0:21.162 |
| ProcessNets-wpgmc | 1:00.007 | 0:47.844 | 0:18.187 | 0:15.353 |

##### *E. Implementation Details of Ensemble Construction, and* ProcessNets-* *and* baseline-* *Configurations*

All empirical analyses, concerning constructions of various network ensembles, and computation of the four network science measures by all seven (five ProcessNets-* and two baseline-*) configurations across these ensembles, were performed on a local server with Intel Xeon platinum 8180 CPU running at 2.5 GHz frequency with 56 physical cores each comprising two (logical) threads, running on a CentOS 7 operating system and 1 TB RAM. All programs were compiled using Python version 3.10 programming language. Python packages NumPy, Pandas, NetworkX, CorALS, and random were primarily used for implementation. For construction of the ensembles alone, parameters governing internal threads such as OMP_NUM_THREADS, MKL_NUM_THREADS, NUMEXPR_NUM_THREADS, OPENBLAS_NUM_THREADS, and VECLIB_MAXIMUM_THREADS were each set to 1 to minimize the memory footprint.

Five logical cores were dedicated to each ensemble using the taskset utility in Linux—each of these cores were assigned to a single configuration of ProcessNets-*; after the ProcessNets-* executions completed, the baseline-* were executed on two of the same cores. This prevented parallel execution of the steps within a configuration across multiple cores, ensuring that each configuration on an ensemble was executed sequentially under identical and controlled conditions, thus enabling fair comparison across the configurations. Since our local server had at least 50 logical cores, we completed the above process for 10 different ensembles in parallel. These different ensembles were generated either from the same simulated dataset but with varied values of $n$, $B$, or $s$, or from different real-world datasets belonging to the same category (see Section II-F).

The parallel execution of different configurations on various ensembles using multiple cores (but with each configuration on an ensemble running sequentially) as described above was used throughout our study. This multi-core benchmarking may lead to less reliable estimates of running times due to potential sharing of hardware resources by different threads. So for two representative ensembles, we disabled the use of multiple cores altogether and ran each configuration on an ensemble exclusively one at a time. This single-core benchmarking actually led to similar performance trends as the multi-core benchmarking (see Table S1). That is, although the absolute running times differ slightly because of the execution environment, the relative ordering and trends in the running times observed among the configurations remain unchanged.

### Appendix B
### Supplementary Results

#### A. *Computation of Closeness Centrality*

We used ensembles of $B = 1000$ networks, each constructed on $n = 100$ nodes and varied sample sizes, from Simulated Ensemble I (see Section II-F2a for details) to analyze performance of the configurations in computation of closeness centrality; MPJS of these ensembles ranged between $[0.67, 0.93]$. Scatter plots depicting the running time taken by different configurations with respect to the sample size is shown in Fig. S1. Clearly, ProcessNets-* show poor running times than baseline-networkx for all values of $s$ (Fig. S1 and "*S2.1.0 Closeness Centrality.xlsx*" in Supplementary Data S2); this result is expected as the incremental dynamic graph algorithm for closeness centrality is dependent on the original graph DS rather than an intermediate DS and does not process the incremental edges as a batch (as discussed in Section II-E in the main text). Since the performance of the framework was subpar even on smaller simulated networks, we did not pursue further analysis on closeness centrality computation.

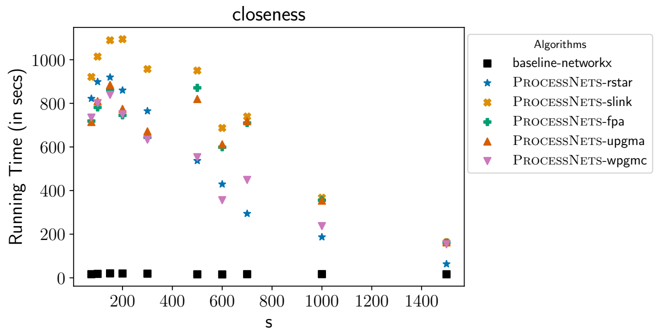

Fig. S1. Running times taken by different configurations to compute closeness centrality in Simulated Ensemble I with $n = 100$, $B = 1000$ and varied $s$.

#### B. *Dissecting* ProcessNets *Speedup using I/O (Input/Output) vs. Processing Times*

Throughout this study, we report the total running times by different configurations to compute the different measures, and calculate the speedup of our framework over a baseline configuration using the ratio of their total running times. To understand the sources of the speedup of our framework such as that of ProcessNets-rstar or ProcessNets-slink, we computed the time required by the configurations to read the input networks from files stored in the hard disk ("input" time), compute the measures ("processing" time), and write the measurements to an output file ("output" time). Note that the input for the baseline-* configurations is the actual networks in the ensemble, whereas the input for the ProcessNets-* configurations is the corresponding NC-tree.

We performed this analysis for category $\mathcal{B}s$ and $\mathcal{S}s$ ensembles on the GTEx Muscle Skeletal tissue on top-1000 highly varying genes, and report the results in Table S2. As the outputs and output times are similar across all configurations, we focus on input and processing times. Our ProcessNets-*, such as ProcessNets-rstar for degree and component size computations and ProcessNets-slink for PR computations, show speedup in both input and processing times compared to the best of the baseline-*. However, the larger of these two speedups pertains to ProcessNets-* performing the input operations much faster than baseline-*, which is a result of NC-trees being a compact data structure to store all the networks in an ensemble (achieving a compression factor of 2.1× or 8.1× for GTEx Muscle Skeletal ensembles Table S2).

#### C. *Dependence of Running Time to Compute PR Centrality on Average Edge Count*

Running times of the different configurations to compute PR centrality in real-world ensembles show some interesting trends. While ProcessNets-* compute the measure faster than the baseline-* in ensembles belonging to category $\mathcal{B}s$, the reverse trend is observed for ensembles in category $\mathcal{S}s$, i.e., the baseline-* have faster running times.

To better understand this discrepancy in the results, we used the gene expression dataset for the Muscle Skeletal tissue with the actual number of genes and varied the FDR cutoff in order to generate ensembles in category $\mathcal{B}s$ with average edge count close to that in ensembles belonging to category $\mathcal{S}s$, and vice-versa. This resulted in four ensembles (two each in categories $\mathcal{B}s$ and $\mathcal{S}s$) grouped into two sets (Table S3):

1) (with Fewer Edges): Here, we reduced the number of edges in the ensemble belonging to category $\mathcal{B}s$ by reducing the FDR cutoff on the adjusted $p$-values to 0.0007, whereas the FDR cutoff used to construct the ensemble in category $\mathcal{S}s$ was fixed at 0.01. This resulted in two ensembles (one from each category) with smaller average edge count as compared to set 2.
2) (with More Edges): Here, we increased the number of edges in the ensemble belonging to category $\mathcal{S}s$ by increasing the FDR cutoff on the adjusted $p$-values to 0.15, whereas the FDR cutoff used to construct the ensemble in category $\mathcal{B}s$ was fixed at 0.01. This resulted in two similar ensembles (one from each category) with higher average edge count relative to set 1.

It may be noted here that MPJS of the ensembles in categories $\mathcal{B}s$ and $\mathcal{S}s$ are 0.2 and 0.6 respectively across both the sets.

Clearly, baseline-self computes PR centralities in the ensembles belonging to set 1 faster than the ProcessNets-*, but the reverse trend occurs for ensembles in set 2, i.e., the ProcessNets-* run faster than the baseline-*. Notably, with increase in the average edge count of an ensemble, the increase in running times of the ProcessNets-* is significantly less relative to the baseline-* (Fig. S2), possibly due to a smaller increase in execution time at each branching point since the new edges are spread across the NC-tree. These results are also in accordance with the trends observed in computation of PR centrality across the real-world ensembles reported in Section IV-C2 and Fig. 5 in the main text.

TABLE S2
BREAKDOWN OF RUNNING TIME IN MUSCLE SKELETAL CATEGORY $\mathcal{B}s$ AND CATEGORY $\mathcal{S}s$ ENSEMBLES WITH TOP-1000 HIGHLY VARYING GENES. HERE 'INPUT', 'PROCESSING', AND 'OUTPUT' REFER TO THE INPUT TIME, PROCESSING TIME AND OUTPUT TIME RESPECTIVELY. NOTE THAT, THE PROCESSING TIME FOR THE `baseline`-NETWORKX CONFIGURATION IS COMPRISED OF TWO PARTS–CREATING THE 'NETWORKX' GRAPH AND THE ACTUAL PROCESSING TIME (SHOWN IN THE SAME ORDER). ALL RUNNING TIMES SHOWN HERE ARE MEASURED IN SECONDS, AND THE BEST VALUE IN EACH ROW IS SHOWN IN BOLD-FACED FONT. NOTE THAT THE REPORTED TOTAL TIME COULD BE SLIGHTLY DIFFERENT FROM THE SUM OF INPUT, PROCESSING, AND OUTPUT TIMES DUE TO INCLUSION OF TIME FOR A FEW EXTRA DRIVER OPERATIONS INCLUDED IN THE TOTAL TIME.

(a) Category $\mathcal{B}s$. $\texttt{MPJS} = 0.44$

| Configurations | | baseline-self | baseline-networkx | ProcessNets-rstar | ProcessNets-slink | ProcessNets-fpa | ProcessNets-upgma | ProcessNets-wpgmc |
|---|---|---|---|---|---|---|---|---|
| Total size to store ensemble | | 172.3 MB | | **81.8 MB** | 112 MB | 83.1 MB | 84.6 MB | 87.3 MB |
| Degree Centrality | Input | 42.14 | 45.14 | **2.58** | 2.77 | 2.69 | 2.64 | 2.65 |
| | Processing | 66.61 | 40.78 + 0.40 | 37.89 | 50.35 | 39.78 | **37.68** | 39.29 |
| | Output | 0.28 | **0.25** | 0.31 | 0.30 | 0.32 | 0.31 | 0.30 |
| | Total Time | 109.50 | 87.00 | 41.43 | 53.89 | 43.46 | **41.10** | 42.73 |
| Component Size | Input | 43.43 | 43.04 | **2.51** | 2.65 | 2.60 | 2.52 | 2.55 |
| | Processing | 19.51 | 36.26 + 11.80 | **13.40** | 17.82 | 14.15 | 13.90 | 14.19 |
| | Output | 0.31 | 0.30 | 0.25 | **0.24** | 0.25 | **0.24** | **0.24** |
| | Total Time | 63.56 | 91.78 | **16.47** | 21.02 | 17.43 | 16.97 | 17.29 |
| PR Centrality | Input | 43.47 | 45.33 | 2.20 | **2.17** | **2.17** | 2.23 | 2.21 |
| | Processing | **18.82** | 34.78 + 39.04 | 36.15 | 32.47 | 36.80 | 36.94 | 37.15 |
| | Output | **0.82** | 0.86 | 0.83 | 0.84 | 0.83 | 0.84 | 0.91 |
| | Total Time | 63.58 | 120.56 | 39.65 | **36.15** | 40.29 | 40.48 | 40.74 |

(b) Category $\mathcal{S}s$. $\texttt{MPJS} = 0.86$

| Configurations | | baseline-self | baseline-networkx | ProcessNets-rstar | ProcessNets-slink | ProcessNets-fpa | ProcessNets-upgma | ProcessNets-wpgmc |
|---|---|---|---|---|---|---|---|---|
| Total size to store ensemble | | 89.6 MB | | **11.1 MB** | 16.8 MB | 16 MB | 16 MB | 13.3 MB |
| Degree Centrality | Input | 22.11 | 23.34 | **1.19** | 1.44 | 1.47 | 1.42 | 1.43 |
| | Processing | 34.61 | 20.94 + 0.36 | **6.44** | 8.80 | 8.33 | 8.48 | 7.21 |
| | Output | **0.29** | 0.30 | 0.31 | 0.34 | 0.31 | 0.38 | 0.34 |
| | Total Time | 57.47 | 45.34 | **8.44** | 11.05 | 10.57 | 10.76 | 9.45 |
| Component Size | Input | 23.76 | 23.31 | **1.21** | 1.42 | 1.45 | 1.44 | 1.42 |
| | Processing | 47.55 | 19.14 + 44.98 | **3.07** | 4.16 | 4.04 | 4.02 | 3.77 |
| | Output | 0.32 | 0.33 | 0.25 | **0.24** | 0.31 | 0.26 | **0.24** |
| | Total Time | 71.93 | 88.14 | **4.84** | 6.13 | 6.10 | 6.02 | 5.74 |
| PR Centrality | Input | 23.06 | 23.82 | **1.25** | 1.46 | 1.48 | 1.47 | 1.46 |
| | Processing | **19.27** | 18.28 + 19.60 | 32.21 | 40.76 | 40.93 | 40.79 | 39.97 |
| | Output | 0.83 | **0.81** | 0.84 | 0.88 | 1.51 | 0.84 | 0.84 |
| | Total Time | 43.63 | 63.07 | **34.78** | 43.58 | 44.40 | 43.58 | 42.74 |

TABLE S3
SUMMARY OF THE FOUR ENSEMBLES USED. HERE "FDR CUTOFF" REPRESENTS THE THRESHOLD USED ON THE ADJUSTED $p$-VALUES DURING CONSTRUCTION OF THE NETWORKS.

| Set Number | Ensembles Constructed using: Bootstrapping (Category $\mathcal{B}s$) | Ensembles Constructed using: Subsampling (Category $\mathcal{S}s$) |
|---|---|---|
| 1 | FDR cutoff $= 0.0007$<br>Avg. EC $= 695812.355$<br>$\texttt{MPJS} = 0.225$ | FDR cutoff $= 0.01$<br>Avg. EC $= 219384.992$<br>$\texttt{MPJS} = 0.593$ |
| 2 | FDR cutoff $= 0.01$<br>Avg. EC $= 3128426.925$<br>$\texttt{MPJS} = 0.235$ | FDR cutoff $= 0.15$<br>Avg. EC $= 2648130.569$<br>$\texttt{MPJS} = 0.560$ |

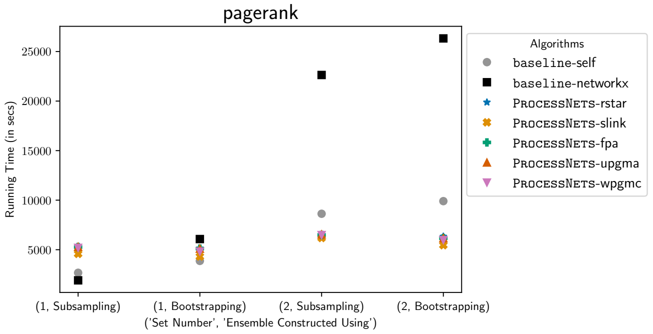

Fig. S2. Runtime of the configurations across the four ensembles mentioned in Table S3 in increasing order of their average edge count.

#### APPENDIX C
#### SUPPLEMENTARY DATA

*Supplementary Data S1:* An example output of results on ensembles constructed using the gene expression dataset on Whole Blood tissue. Link: https://drive.google.com/drive/folders/1PMPtwPyJEzOKTK-3NfIVBg6wgjbNboR4?usp=sharing.

*Supplementary Data S2:* Running times of the ProcessNets-* and `baseline-*` configurations for different ensembles. Link: https://drive.google.com/drive/folders/1RwzqlyXvdUoVBRQewmQRRWgzldrkces9?usp=sharing. The files included in this folder are structured as follows:

- S2.1 Simulated Ensemble I: Contains running times on ensembles from Simulated Ensemble I
  - S2.1.0 Closeness Centrality.xlsx: results for computing closeness centrality
  - S2.1.1 n,B effect with fixed s.xlsx: running times when $n$ and $B$ are varied with fixed $s = 500$ (data underlying Fig. 2A)
  - S2.1.2 n,s effect with fixed B.xlsx: running times when $n$ and $s$ are varied with fixed $B = 1000$ (data underlying Fig. 2B)
  - S2.1.3 A folder containing interactive 3D plots corresponding to Fig. 2A-B
- S2.2 Simulated Ensemble II.xlsx: Contains running times on all ensembles from Simulated Ensemble II (Fig. 2C)
- S2.3 Real World Ensembles: Contains running times on all real-world ensembles across tissues
  - S2.3.1 Muscle Skeletal: contains results on ensembles from Muscle Skeletal tissue (data underlying Fig. 3A-C) – one file for each category
    - S2.3.1A Category Bs.xlsx
    - S2.3.1B Category Ss.xlsx
    - S2.3.1C Category Ssf.xlsx
  - S2.3.2 All Chosen Tissues: contains results on ensembles from all other tissues (data underlying Fig. 4A-B) – one file for each category
    - S2.3.2A Category Bs.xlsx
    - S2.3.2B Category Ss.xlsx

*Supplementary Data S3:* Average Ranking of the ProcessNets-* and `baseline-*` configurations for different ensembles. Link: https://docs.google.com/spreadsheets/d/1qwDdnbN92-F8yeVtFn7brHw4UFZMBAUK/edit?usp=sharing&ouid=101900263397605157998&rtpof=true&sd=true.

#### REFERENCES

[1] J. Leskovec, A. Rajaraman, and J. D. Ullman, *Mining of Massive Datasets*, 3rd ed. Cambridge University Press, 2020.

[2] T. H. Cormen, C. E. Leiserson, R. L. Rivest, and C. Stein, *Introduction to Algorithms*, 3rd ed. The MIT Press, 2009.

[3] J. Riedy, "Updating PageRank for Streaming Graphs," in *2016 IEEE International Parallel and Distributed Processing Symposium Workshops (IPDPSW)*. IEEE, 2016, pp. 877–884.

[4] E. Nathan and D. A. Bader, "Incrementally updating Katz centrality in dynamic graphs," *Social Network Analysis and Mining*, vol. 8, no. 26, 2018.

[5] Newman, Mark, *Networks*, 2nd ed. Oxford University Press, 2018.

[6] C. C. Paige and M. A. Saunders, "Solution of Sparse Indefinite Systems of Linear Equations," *SIAM Journal on Numerical Analysis*, vol. 12, no. 4, pp. 617–629, 1975.

[7] L. C. Freeman, "Centrality in social networks conceptual clarification," *Social Networks*, vol. 1, no. 3, pp. 215–239, 1978.

[8] A. E. Sariyüce, K. Kaya, E. Saule, and Ü. V. Çatalyiirek, "Incremental algorithms for closeness centrality," in *2013 IEEE International Conference on Big Data*. IEEE, 2013, pp. 487–492.

[9] P. Langfelder and S. Horvath, "WGCNA: an R package for weighted correlation network analysis," *BMC Bioinformatics*, vol. 9, no. 1, 2008.

[10] G. G. Lemoine, M. P. Scott-Boyer, B. Ambroise, O. Périn, and A. Droit, "GWENA: gene co-expression networks analysis and extended modules characterization in a single Bioconductor package," *BMC Bioinformatics*, 2021.

[11] M. A. Cortes-Guzman and V. Treviño, "CoGTEx: Unscaled system-level coexpression estimation from GTEx data forecast novel functional gene partners," *PLOS One*, 2024.

[12] P. Raina, R. Guinea, K. Chatsirisupachai, I. Lopes, Z. Farooq, C. Guinea, C. A. Solyom, and J. P. D. Magalhães, "GeneFriends: gene co-expression databases and tools for humans and model organisms," *Nucleic Acids Research*, 2023.

[13] S. Mahapatra, N. A. Subramanian, and M. Narayanan, "Uncertainty-Aware Gene Rankings Reveal Key Players in Coexpression Networks," *bioRxiv*, 2025.
